# DNAM-1 immunoreceptor integrates innate and adaptive immune programs to drive intestinal inflammation

**DOI:** 10.64898/2026.03.30.715436

**Authors:** Natsuki Ide, Kazuki Sato, Kyoko Oh-oka, Mariana Silva Almeida, Fumie Abe, Tae-hyeong Kim, Chigusa Nakahashi-Oda, Kazuko Shibuya, Akira Shibuya

## Abstract

Innate and adaptive immune responses play critical roles in the pathogenesis of inflammatory bowel disease (IBD), yet the molecular pathways integrating these responses remain elusive. Here, we identify DNAM-1 immunoreceptor as a central driver of colitis through distinct, cell type–specific mechanisms. Transcriptomic analyses of human and murine group 3 innate lymphoid cells (ILC3s) revealed DNAM-1 as a conserved IL-23–responsive surface molecule associated with inflammatory cytokine production. In an innate immune–driven anti-CD40 monoclonal antibody (mAb)–induced colitis model, DNAM-1 expressed on ILC3s promoted intestinal inflammation by enhancing IL-22 and GM-CSF production via the integration of the Akt–mTORC1–HIF-1α signaling pathway. Genetic ablation or antibody-mediated blockade of DNAM-1 attenuated inflammatory cytokine production and disease severity. Paradoxically, in T cell–dependent colitis, DNAM-1 expression on dendritic cells, but not on ILC3s or CD4⁺ T cells, exacerbated disease by promoting dendritic cell activation and pathogenic Th1 and Th17 differentiation. Notably, therapeutic blockade of DNAM-1 ameliorated disease in both colitis models and exerted complementary effects when combined with anti-TNF therapy, accompanied by modulation of immune activation programs distinct from those regulated by TNF inhibition. Collectively, these findings establish DNAM-1 as a pivotal regulator of intestinal inflammation bridging innate and adaptive immunity and identify DNAM-1 blockade as a next-generation therapeutic strategy for IBD.

**Highlight:**

► DNAM-1 is an IL-23-responsive receptor conserved in human and mouse ILC3s.
► DNAM-1 on ILC3s drives innate colitis via Akt-mTORC1-HIF-1α signaling.
► DNAM-1 on DCs promotes T cell-dependent colitis by inducing Th1/Th17 cells.
► DNAM-1 blockade targets immune pathways distinct from TNF inhibition.
► Combined DNAM-1 and TNF blockade shows additive therapeutic efficacy in colitis.

## Introduction

Inflammatory bowel disease (IBD), comprising Crohn’s disease (CD) and ulcerative colitis (UC), is a group of chronic relapsing inflammatory disorders of the gastrointestinal tract driven by complex interactions among genetic susceptibility, intestinal microbiota, mucosal immune responses, and environmental factors ^1–3^. Accumulating evidence indicates that innate immune dysregulation acts as an upstream determinant of disease initiation, as defects in epithelial barrier function, innate immune sensing, autophagy, and activation of innate immune cells disrupt mucosal homeostasis and promote aberrant inflammatory responses to commensal microbes ^4^. In addition, adaptive immune responses—particularly pathogenic CD4⁺ T cell differentiation—are critical for the perpetuation of chronic intestinal inflammation ^5^. Although biologic therapies targeting TNF-α, IL-12/IL-23, and α4β7 integrin have substantially improved disease outcomes, a considerable proportion of patients remain refractory or lose response over time ^6^, underscoring the need to identify additional pathogenic pathways and therapeutic targets.

Murine models of IBD have been instrumental in dissecting the cellular and molecular mechanisms underlying IBD pathogenesis. Among these, the anti-CD40 antibody–induced colitis model and the CD4⁺ T cell transfer model represent complementary systems that recapitulate distinct aspects of disease. The anti-CD40 antibody–induced colitis model captures rapid, innate immune–driven inflammation, where activation of CD40 on antigen-presenting cells induces group 3 innate lymphoid cells (ILC3s)–dependent pathology independent of T cells ^7^. ILC3s are selectively enriched in the intestinal mucosa and play central roles in maintaining epithelial barrier integrity and host–microbiota homeostasis ^8, 9^. ILC3s rapidly respond to environmental cues and contribute to intestinal inflammation through the production of effector cytokines implicated in IBD pathogenesis, including IL-22, IL-17A, IFN-γ, and GM-CSF ^10–13^. While IL-22 is classically associated with epithelial repair and barrier protection, excessive or context-dependent IL-22 signaling can exacerbate tissue inflammation ^14–16^. Similarly, ILC3-derived GM-CSF promotes recruitment and activation of neutrophils and inflammatory monocytes, amplifying intestinal pathology ^14, 17^. In contrast, the CD4⁺CD45RB^high^ T cell transfer model recapitulates chronic, microbiota-dependent colitis driven by uncontrolled Th1/Th17 responses in the absence of regulatory T cells ^18, 19^. Together, these complementary models reflect key features of human IBD, spanning early innate immune activation to sustained adaptive immune–mediated inflammation, and provide a framework for mechanistic and therapeutic studies.

DNAM-1 (*CD226*) is an immunoglobulin-like receptor expressed on a broad range of innate and adaptive immune cells, including NK cells, cytotoxic and helper T cells, dendritic cells, and subsets of innate lymphoid cells ^20, 21^. Engagement of DNAM-1 with its ligands CD155 (PVR) and CD112 (nectin-2) delivers activating signals that enhance cytokine production and effector function ^22, 23^. DNAM-1 signaling is counterbalanced by inhibitory receptors such as TIGIT and CD96, which compete for shared ligands and fine-tune immune activation thresholds ^24, 25^. While DNAM-1 has been extensively studied in cytotoxic lymphocytes and cancer immunity, its role in mucosal immune regulation remains poorly understood. Notably, dysregulation of the DNAM-1–ligand axis has been linked to inflammatory diseases, including pediatric IBD ^26^, suggesting that DNAM-1 may function at the interface of innate and adaptive immunity in the gut. However, whether DNAM-1 regulates both innate immune–driven and T cell–dependent intestinal inflammation has not been directly addressed.

In this study, we investigated the role of DNAM-1 in intestinal inflammation using complementary models of innate and adaptive immune–driven colitis. We demonstrate that DNAM-1 promotes ILC3-mediated intestinal inflammation in an anti-CD40 mAb–induced colitis model by enhancing IL-22 and GM-CSF production through activation of the Akt–mTORC1–HIF-1α signaling pathway. In contrast, in a T cell transfer colitis model, DNAM-1 expressed on dendritic cells, rather than on T cells or ILC3s, exacerbates disease by promoting pathogenic Th1 and Th17 differentiation. Furthermore, therapeutic blockade of DNAM-1 ameliorates intestinal inflammation in both models and exhibits additive effects when combined with anti-TNF therapy. Together, these findings identify DNAM-1 as a previously unrecognized regulator of intestinal inflammation bridging innate and adaptive immunity.

## Results

### DNAM-1 on ILC3s associates with intestinal inflammation in humans and mice

IL-23 activates ILC3s and is elevated in patients with inflammatory bowel disease (IBD) ^10^. To investigate molecular pathways by which ILC3s contribute to human intestinal inflammation, we analyzed a public microarray dataset of human tonsillar ILC3s before and after ex vivo stimulation with IL-23 in combination with IL-1β and IL-15 (GSE132843) ^27^.

This analysis identified 567 genes that were upregulated following stimulation (Figure 1A). Of these, 29 genes were located within established IBD risk loci ^28^. To identify surface molecules potentially linking ILC3 activation to cytokine production, we interrogated gene ontology (GO) categories associated with cytokine production (GO:0001816) and cell surface molecules (GO:0009986). This approach yielded two candidate genes, *CD226* and *SLAMF1*, that fulfilled all selection criteria (Figure 1A).

**Figure 1.**
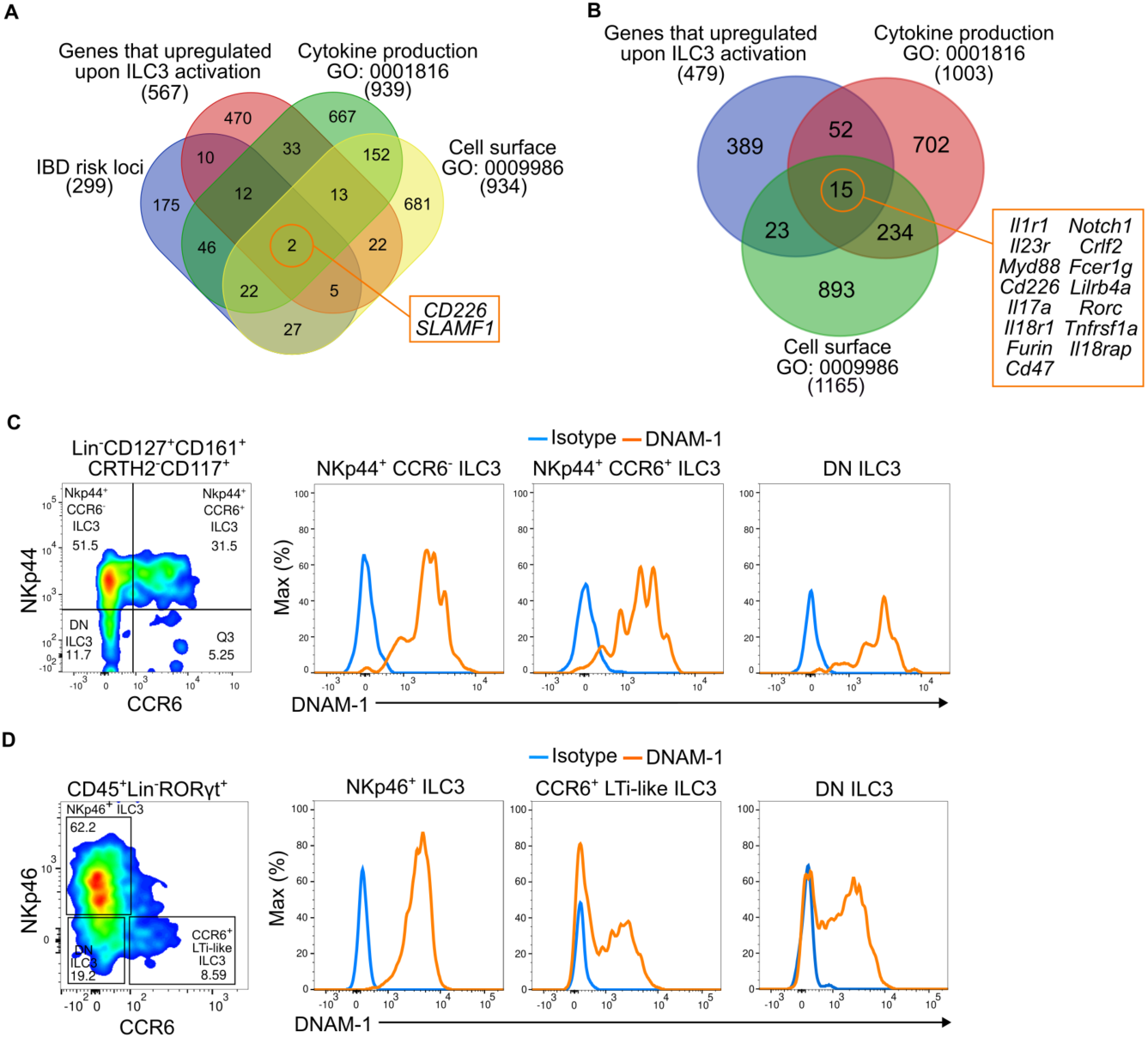
DNAM-1 on ILC3s associates with intestinal inflammation in humans and mice. **(A, B)** Venn diagrams showing the overlap among genes upregulated upon ILC3 activation, IBD risk loci (A only), genes associated with cytokine production (GO:0001816), and cell surface molecules (GO:0009986) in human tonsillar ILC3s stimulated with IL-23, IL-1β, and IL-15 (GSE132843) (A) and mouse small intestinal ILC3s stimulated with IL-23 (GSE229976) (B). Numbers indicate the count of genes in each intersection. Genes listed in (B) represent the intersection of cytokine production and cell surface categories upregulated upon stimulation. **(C, D)** Representative flow cytometric analysis of DNAM-1 expression (orange) versus isotype control (blue) on ILC3 subsets from healthy human peripheral blood (C) and the intestine of naïve SCID mice (D). Subsets were gated on Lin⁻ CD127⁺ CD161⁺ CRTH2⁻ CD117⁺ ILC3s (C) and CD45⁺ Lin⁻ RORγt⁺ cells (D) and defined as NKp44⁺ CCR6⁻, NKp44⁺ CCR6⁺, and NKp44⁻ CCR6⁻ subsets in humans (C), and as NKp46⁺ ILC3s, CCR6⁺ LTi-like ILC3s, and NKp46⁻ CCR6⁻ double-negative (DN) ILC3s in mice (D).

To assess conservation across species, we analyzed a single-cell RNA-sequencing dataset of mouse small intestinal ILC3s stimulated ex vivo with IL-23 (GSE229976) ^29^. Differential expression analysis combined with the same GO categories identified 15 genes encoding cell surface molecules, including *Cd226*, that were upregulated upon IL-23 stimulation and associated with cytokine production in murine intestinal ILC3s (Figure 1B). Among these candidates, DNAM-1, encoded by *CD226* in humans and *Cd226* in mice, was the only gene shared between the human and mouse datasets.

We then examined DNAM-1 expression on ILC3s in the peripheral blood from healthy volunteers. Human ILC3s were identified by flow cytometry as Lin- (CD138, FcεRIα, CD3, CD11b, CD34, CD56, CD14, and CD19) CD127+CD161+CD117+ CRTH2- cells (Supplemental Figure 1A). The ILC3s were further divided into NKp44+ CCR6-, NKp44+ CCR6+, and NKp44- CCR6- subpopulations. All these subpopulations similarly expressed DNAM-1 (Figure 1C). To investigate the role of DNAM-1 on ILC3s in intestinal inflammation in vivo, we utilized an anti-CD40 mAb–induced colitis model in SCID mice, in which IL-23 produced by anti-CD40 mAb–stimulated myeloid cells activates ILC3s and drives intestinal pathology ^7, 30^. Flow cytometric analysis demonstrated high DNAM-1 expression on Lin⁻ (CD11b, CD11c, CD49b, KLRG1) NKp46⁺ CCR6⁻ ILC3s and Lin⁻ NKp46⁻ CCR6⁻ ILC3s, which represent the predominant ILC3 subsets in the intestines of SCID mice (Figure 1D; Supplemental Figure 1B). DNAM-1 expression was also detected on NK cells, ILC1s, ILC2s, and dendritic cells, whereas expression was minimal on macrophages, neutrophils, monocytes, and eosinophils (Supplemental Figures 1B, C).

### DNAM-1 blockade ameliorates anti-CD40 mAb–induced intestinal inflammation

To assess the functional contribution of DNAM-1 to intestinal inflammation, SCID mice were intraperitoneally administered either a neutralizing anti–DNAM-1 mAb or a control mAb together with anti-CD40 mAb on day 0, and disease severity was evaluated on day 5. Anti–DNAM-1 mAb treatment resulted in reduced colonic inflammation, as indicated by lower histological scores and decreased colon weight–to–length ratios (Figures 2A, B).

**Figure 2.**
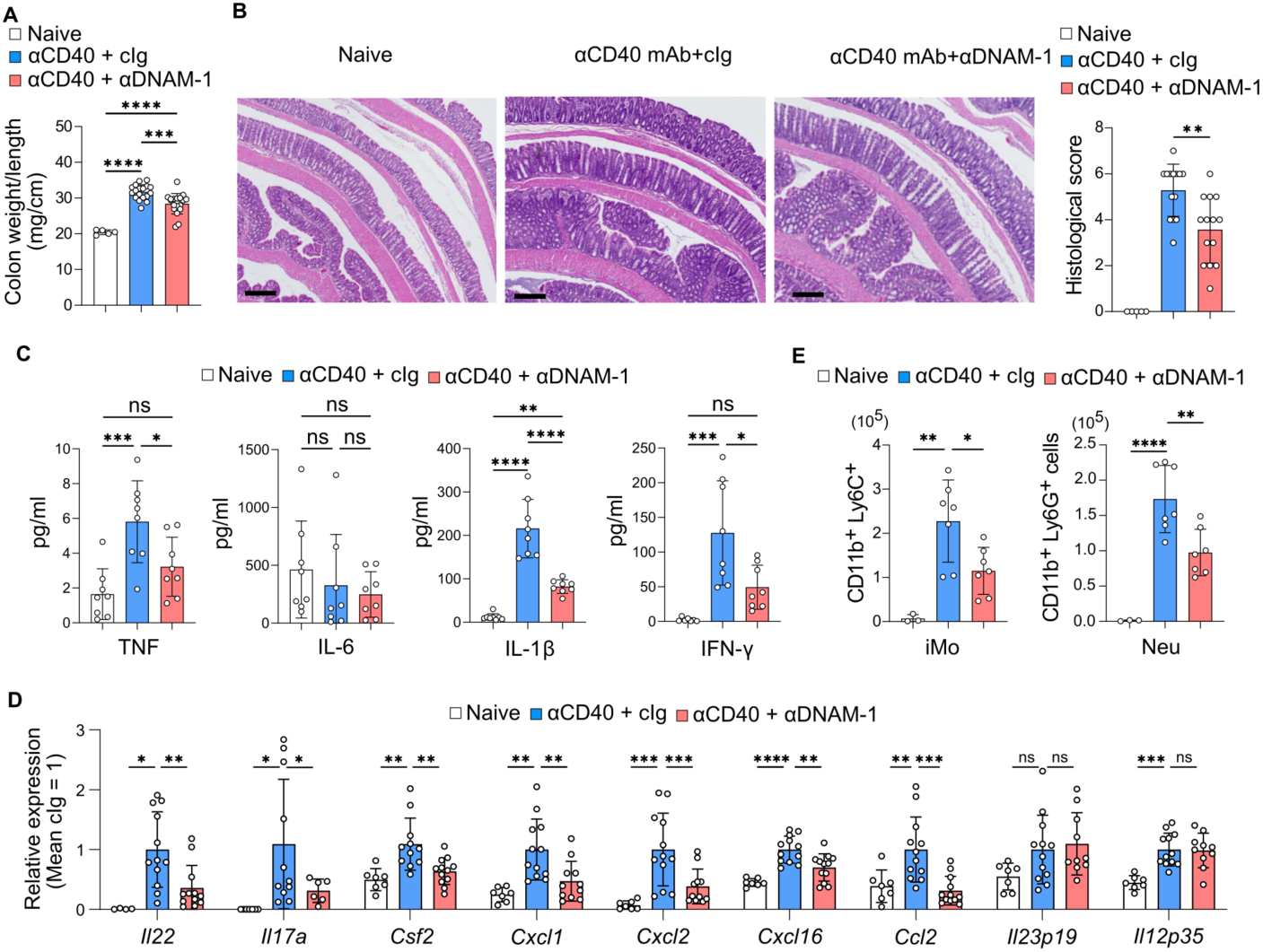
DNAM-1 blockade ameliorates anti-CD40 mAb–induced intestinal inflammation. **(A, B)** Colon weight-to-length ratio (A) and representative H&E-stained colon sections and histological scores (B) from naive mice (n = 5) and mice treated with control Ig (cIg) or anti–DNAM-1 mAb (n = 14) at day 5 post anti-CD40 mAb injection. Scale bars, 100 μm. **(C, D)** Cytokine concentrations in colon tissue culture supernatants (n = 8 in each group) (C) and relative mRNA expression of inflammatory cytokine genes in the colon from naïve mice (n = 7) and mice treated with cIg (n = 12) or anti–DNAM-1 mAb (n = 12) (D) at day 5 post-anti-CD40 mAb. Values in (D) are normalized to the cIg group. (E) Absolute numbers of CD11b⁺ Ly6C⁺ inflammatory monocytes (iMo) and CD11b⁺ Ly6G⁺ neutrophils (Neu) in the colon from naive mice (n = 3) and mice treated with cIg or anti–DNAM-1 mAb (n = 7) at day 3 after anti-CD40 mAb injection. Data in (A–D) were pooled from two independent experiments; **(E)** from three. Data represent mean ± SD. Statistical significance was determined by one-way ANOVA with Tukey’s post-hoc test; *p < 0.05, **p < 0.01, ***p < 0.001, ****p < 0.0001; ns, not significant.

Consistent with these findings, body weight loss was attenuated in anti–DNAM-1 mAb–treated mice compared with control mAb–treated mice (Supplemental Figure 2A).

Levels of inflammatory cytokines, including TNF, IL-1β, and IFN-γ, were reduced in colon tissue culture supernatants from anti–DNAM-1 mAb–treated mice (Figure 2C). In addition, colonic expression of inflammatory cytokines (*Il22, Il17a,* and *Csf2*) and chemokines (*Cxcl1, Cxcl2, Cxcl16,* and *Ccl2*) was significantly decreased following DNAM-1 blockade (Figure 2D). In contrast, expression of Il23p19 and Il12p35 did not differ between cIg- and anti–DNAM-1 mAb–treated groups. DNAM-1 neutralization was also associated with reduced infiltration of neutrophils and inflammatory monocytes into the colon (Figure 2E; Supplemental Figure 2B). Together, these data demonstrate that DNAM-1 contributes to anti-CD40 mAb–induced intestinal inflammation and that its blockade attenuates disease severity.

### DNAM-1 promotes intestinal inflammation through inflammatory cytokine production by ILC3s

We next examined the contribution of DNAM-1 expressed on ILC3s to the pathogenesis of anti-CD40 mAb–induced intestinal inflammation. Flow cytometric analysis demonstrated marked upregulation of intracellular inflammatory cytokines, including IL-22, IL-17A, GM-CSF, and IFN-γ, in intestinal ILC3s on day **3** following anti-CD40 mAb administration (Figure 3A; Supplemental Figure 3A). Blockade of DNAM-1 with a neutralizing anti–DNAM-1 mAb significantly reduced the production of all these cytokines (Figure 3A), indicating that DNAM-1–mediated signaling promotes inflammatory cytokine production by ILC3s during anti-CD40 mAb–induced colitis.

**Figure 3.**
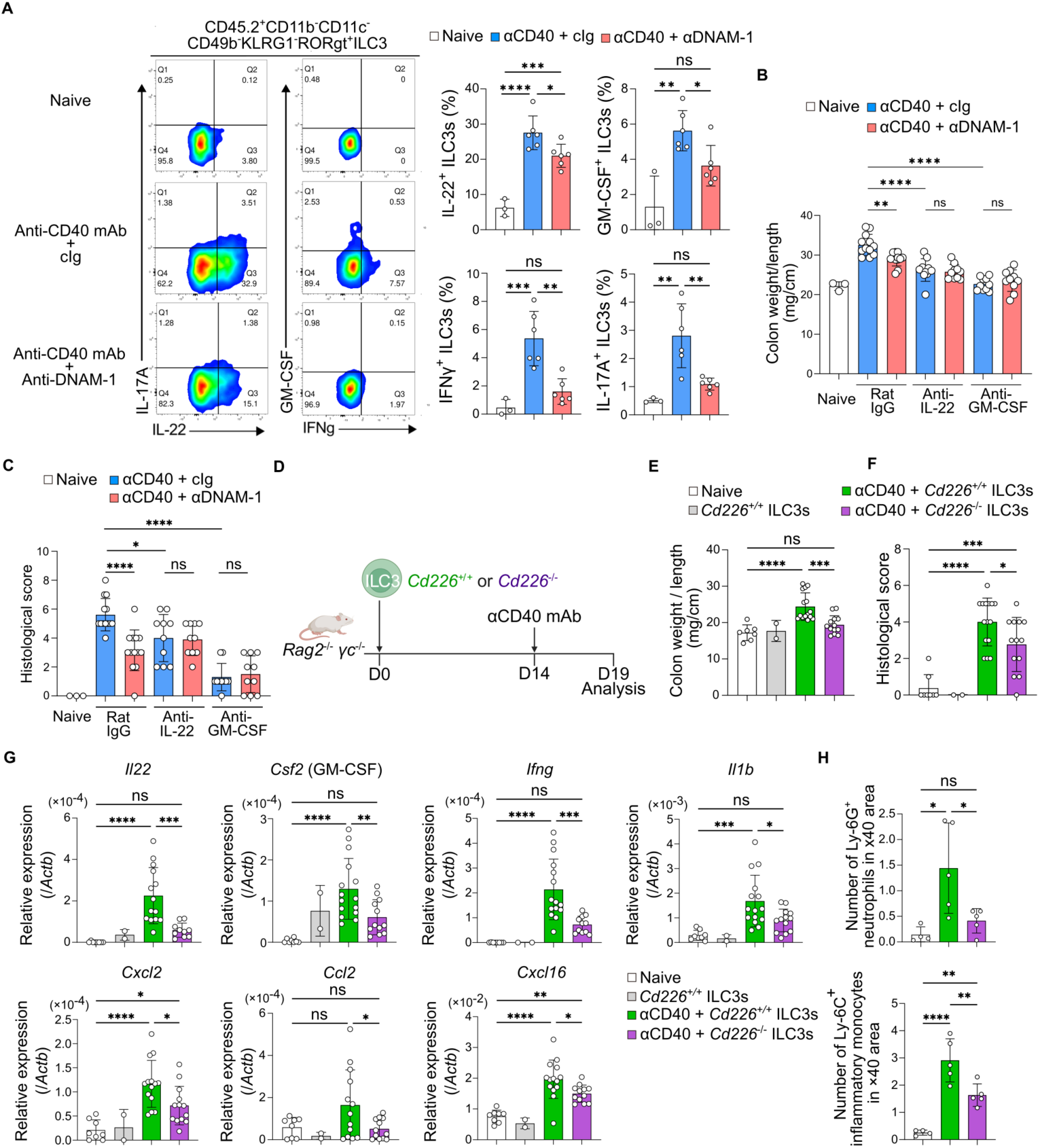
DNAM-1 promotes intestinal inflammation through inflammatory cytokine production by ILC3s. 1. Representative flow cytometry plots (left) and frequencies (right) of IL-22⁺, GM-CSF⁺, IL-17A⁺, and IFN-γ⁺ cells among intestinal ILC3s (CD45.2⁺ CD11b⁻ CD11c⁻ CD49b⁻ KLRG1⁻ RORγt⁺) from naïve mice (n = 3) and mice treated with cIg or anti–DNAM-1 mAb (n = 6) at day 3 after anti-CD40 mAb injection. **(B, C)** Colon weight-to-length ratio (B) and histological scores (C) from naïve mice (n = 3) and mice co-treated with anti-CD40 mAb plus cIg or anti–DNAM-1 mAb, in combination with rat IgG, anti–IL-22 mAb, or anti–GM-CSF mAb (n = 5–6 per group) at day 5 post-injection. Treatment with anti–IL-22 or anti–GM-CSF mAb reduced disease severity to comparable levels irrespective of concomitant DNAM-1 blockade. (D) Schematic illustration of the experimental design of the adoptive transfer experiment. Intestinal ILC3s from *Cd226*⁺^/^⁺ or *Cd226*⁻^/^⁻ mice were transferred into Rag2⁻^/^⁻ γc⁻^/^⁻ mice on day 0, followed by anti-CD40 mAb injection on day 14 and analysis on day 19. **(E–G)** Colon weight-to-length ratio (E), histological scores (F), and relative mRNA expression of indicated cytokines and chemokines (normalized to Actb) (G) in the colon from naïve mice (n = 8), mice reconstituted with Cd226⁺^/^⁺ ILC3s without colitis induction (n = 2), and mice reconstituted with Cd226⁺^/^⁺ (n = 15) or Cd226⁻^/^⁻ (n = 13) ILC3s at day 5 after anti-CD40 mAb injection. RNA in (G) was extracted from formalin-fixed paraffin-embedded (FFPE) colon sections. **(H)** Quantification of Ly-6G⁺ neutrophils and Ly-6C⁺ inflammatory monocytes per ×40 field in immunohistochemically stained colon sections from naïve mice (n = 4) and mice reconstituted with Cd226⁺/⁺ or Cd226⁻/⁻ ILC3s (n = 5) at day 5 after anti-CD40 mAb injection. Data in (A–C) and (H) are representative of or pooled from two independent experiments; data in (E–G) were pooled from four independent experiments. Data represent mean ± SD. Statistical significance was determined by one-way ANOVA with Tukey’s post-hoc test; *p < 0.05, **p < 0.01, ***p < 0.001, ****p < 0.0001; ns, not significant.

To assess the functional relevance of ILC3-derived cytokines in this model, we administered neutralizing mAbs against IL-22 or GM-CSF together with either anti–DNAM-1 mAb or control mAb to anti-CD40 mAb–treated mice. Treatment with either anti–IL-22 or anti–GM-CSF mAb in combination with control mAb reduced colon weight–to–length ratios and histological scores to levels comparable to those observed in mice treated with the corresponding cytokine-neutralizing mAb together with anti–DNAM-1 mAb (Figures 3B, C; Supplemental Figure 3B). These findings indicate that DNAM-1 promotes the development of anti-CD40 mAb–induced colitis at least in part through induction of IL-22 and GM-CSF production by ILC3s.

To directly determine the contribution of DNAM-1 expressed on ILC3s to disease exacerbation, we performed adoptive transfer experiments using *Rag2*⁻/⁻γc⁻/⁻ mice, which lack all lymphocytes, including ILCs. These mice were reconstituted with intestinal ILC3s isolated from wild-type (WT) or DNAM-1–deficient (*Cd226⁻/⁻*) mice and subsequently administered anti-CD40 mAb to induce colitis 14 days after ILC3 transfer (Figure 3D). Flow cytometric analysis confirmed comparable reconstitution of WT and *Cd226⁻/⁻* ILC3s in the colon (Supplemental Figure 3C). Mice receiving WT ILC3s exhibited increased colon weight–to–length ratios (Figure 3E) and higher histological scores (Figure 3F; Supplemental Figure 3D) compared with mice reconstituted with *Cd226⁻/⁻* ILC3s.

Consistent with disease severity, expression levels of genes encoding proinflammatory cytokines and chemokines were elevated in colonic tissues of mice receiving WT ILC3s relative to those receiving *Cd226⁻/⁻* ILC3s (Figure 3G). Notably, expression of certain inflammatory genes, including *Tnfa, Il17a,* and *Cxcl1*, did not differ between groups (Supplemental Figure 3E), in contrast to the broader suppression observed following systemic DNAM-1 blockade (Figures 2C and 2D), suggesting that DNAM-1 expressed on cell types other than ILC3s may also contribute to the inflammatory response. In line with enhanced chemokine expression, immunohistochemical analysis revealed increased infiltration of Ly-6G⁺ neutrophils and Ly-6C⁺ inflammatory monocytes in the colons of mice reconstituted with WT ILC3s (Figure 3H; Supplemental Figures 3F, G). Collectively, these results support a critical role for DNAM-1 expressed on ILC3s in promoting anti-CD40 mAb–induced intestinal inflammation through the induction of inflammatory cytokine production.

### DNAM-1 promotes IL-22 production in ILC3s through Akt/mTORC1/HIF-1α signaling

Administration of anti-CD40 mAb activates intestinal myeloid cells and induces IL-23 production, thereby promoting intestinal inflammation through activation of ILC3s in SCID mice ^7^. To investigate the mechanisms by which DNAM-1 expressed on ILC3s exacerbates anti-CD40 mAb–induced colitis, we established a coculture system consisting of intestinal ILC3s isolated from naïve *Rag*⁻/⁻ or *Rag*⁻/⁻ *Cd226*⁻/⁻ mice and activated CD11b⁺/CD11c⁺ myeloid cells derived from anti-CD40 mAb–treated *Rag*⁻/⁻ mice (Figure 4A; Supplemental Figure 4A).

**Figure 4.**
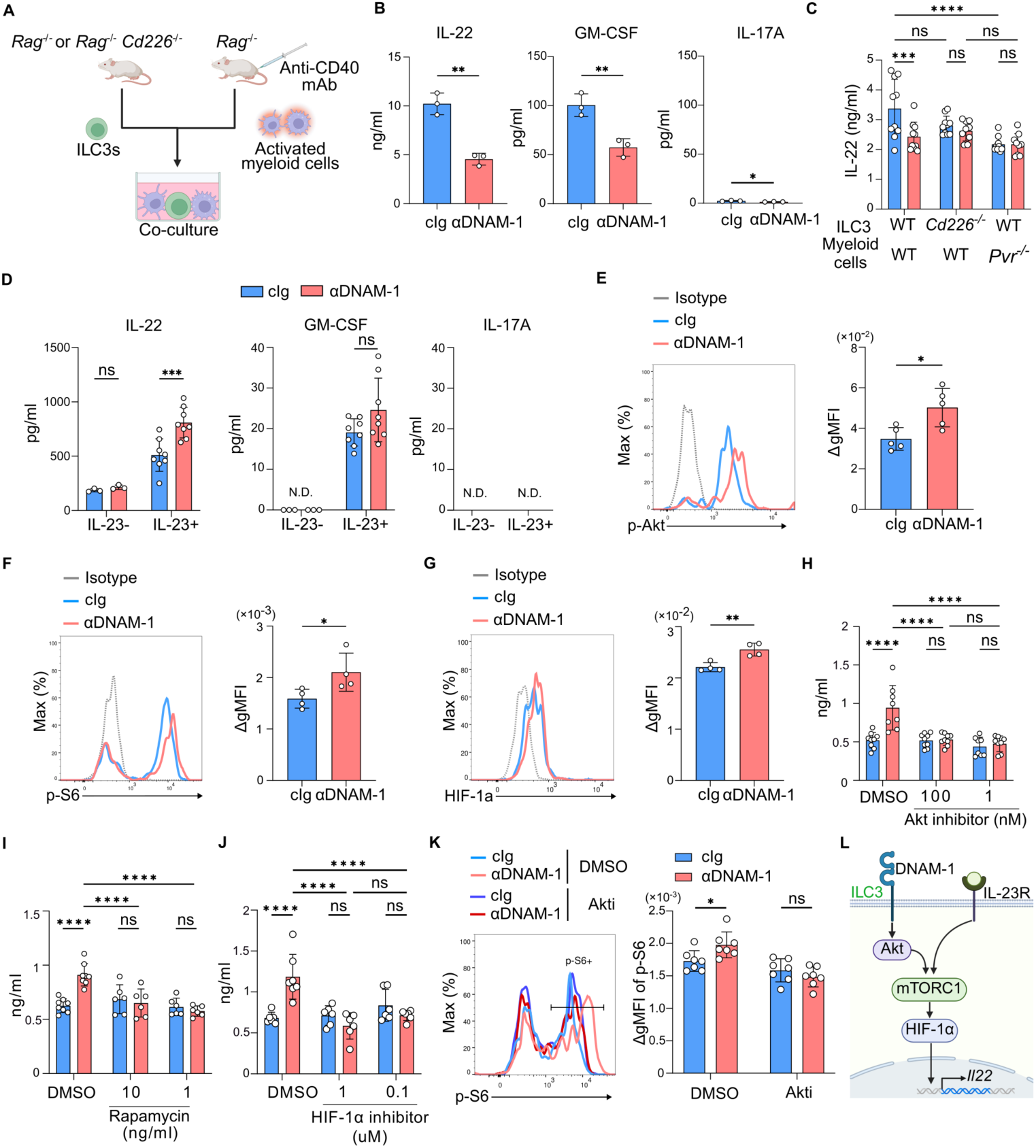
DNAM-1 promotes IL-22 production in ILC3s via the Akt–mTORC1–HIF-1α signaling pathway. 1. Experimental setup for coculture of intestinal ILC3s (from naïve mice) and activated CD11b+ myeloid cells (from anti-CD40-treated mice). **(B)** IL-22, GM-CSF, and IL-17A levels in coculture supernatants treated with cIg (n = 3) or anti–DNAM-1 mAb (n = 3). **(C)** IL-22 production in cocultures using combinations of WT or *Cd226^-/-^*ILC3s and WT or *Pvr^-/-^* myeloid cells. **(D)** IL-22, GM-CSF, and IL-17A production by purified ILC3s stimulated with plate-bound anti–DNAM-1 mAb and/or IL-23 for 24 h. **(E**--**G)** Representative histograms and MFI of phospho-Akt (E), phospho-S6 (F), and HIF-1α (G) in ILC3s stimulated with plate-bound cIg (n = 4-5) or anti–DNAM-1 (n = 4-5) mAbs. **(H**--**J)** IL-22 production by plate-bound cIg (n = 6-9) or anti–DNAM-1 (n = 6-8) mAb-stimulated ILC3s in the presence or absence of inhibitors for Akt (H), mTORC1 (I), or HIF-1α (J). **(K)** Phospho-S6 levels in DNAM-1-stimulated ILC3s with or without Akt inhibitor. **(L)** Schematic model of the DNAM-1–Akt–mTORC1–HIF-1α signaling axis in ILC3s. Data are representative of at least three independent experiments. Data represent mean ± SD. Statistical significance was determined by one-way ANOVA with Tukey’s post-hoc test.

Flow cytometric analysis revealed that multiple myeloid subsets, including macrophages, dendritic cells, and inflammatory monocytes, expressed the DNAM-1 ligand CD155, whereas CD112 expression was minimal (Supplemental Figure 4B). These findings suggest that DNAM-1 on ILC3s may deliver an activating signal through interaction with CD155 on myeloid cells. Consistent with this model, DNAM-1 blockade using a neutralizing anti–DNAM-1 mAb significantly reduced the production of IL-22, GM-CSF, and IL-17A in cocultures of ILC3s and activated myeloid cells (Figure 4B). Similarly, coculture with either *Cd226^⁻/⁻^* ILC3s or *Pvr^⁻/⁻^* myeloid cells resulted in reduced IL-22 production to levels comparable to those observed following DNAM-1 blockade (Figure 4C), indicating that DNAM-1–CD155 interactions enhance IL-22 production by ILC3s.

To determine whether DNAM-1 directly regulates cytokine production in ILC3s, purified intestinal ILC3s were stimulated with plate-bound anti–DNAM-1 mAb in the presence or absence of IL-23. In the absence of IL-23, ILC3s produced little or no IL-22, GM-CSF, or IL-17A (Figure 4D). IL-23 stimulation induced IL-22 and GM-CSF production, but not IL-17A, whereas combined stimulation with IL-23 and DNAM-1 further enhanced IL-22 production (Figure 4D). These data indicate that DNAM-1 cooperates with IL-23 to selectively augment IL-22 production by intestinal ILC3s.

Previous studies have demonstrated that DNAM-1 signals through the PI3K–Akt pathway in NK cells and ILC2s ^23, 31^. Consistently, cross-linking DNAM-1 with plate-bound anti–DNAM-1 mAb increased Akt phosphorylation in ILC3s (Figure 4E). DNAM-1 stimulation also enhanced phosphorylation of S6 (Figure 4F), a downstream target of mTORC1 implicated in ILC3 effector function, as well as expression of HIF-1α (Figure 4G), a transcription factor downstream of mTORC1 known to promote IL-22 production. Pharmacological inhibition of Akt, mTORC1/S6, or HIF-1α reduced IL-22 production to levels comparable to those observed in unstimulated ILC3s (Figure 4H–J). Moreover, Akt inhibition suppressed S6 phosphorylation following DNAM-1 stimulation (Figure 4K). Collectively, these findings identify a DNAM-1–dependent signaling pathway in ILC3s that promotes IL-22 production through activation of the Akt/mTORC1/HIF-1α axis (Figure 4L).

### Anti–DNAM-1 mAb treatment ameliorates T cell–dependent intestinal inflammation

To extend our analysis of DNAM-1 in IBD, we employed a T cell–dependent colitis model ^4, 32, 33^. Naïve CD4⁺CD25⁻CD45RBhigh T cells from BALB/c mice were adoptively transferred into SCID mice on day 0 (Figure 5A; Supplemental Figure 5A). Recipient mice received weekly intraperitoneal injections of either anti–DNAM-1 or control mAb from day 0 until one week prior to analysis. Although body weight loss was comparable between groups (Supplemental Figure 5B), mice treated with anti–DNAM-1 mAb exhibited improved histopathological features in the colon, including reduced colon weight-to-length ratios and attenuation of epithelial damage and inflammatory cell infiltration (Figures 5B, C). Since DNAM-1 is expressed on CD4⁺ T cells, we next tested whether DNAM-1 on transferred T cells contributes to disease. Naïve CD4⁺ T cells from WT or *Cd226*⁻/⁻ mice were adoptively transferred into SCID recipients (Figure 5D). Unexpectedly, both groups developed similar histopathology, as indicated by histological scores and colon weight-to-length ratios on day 49 (Figures 5E, F; Supplemental Figures 5C, D), suggesting that DNAM-1 on recipient immune cells rather than on transferred T cells drives disease.

**Figure 5.**
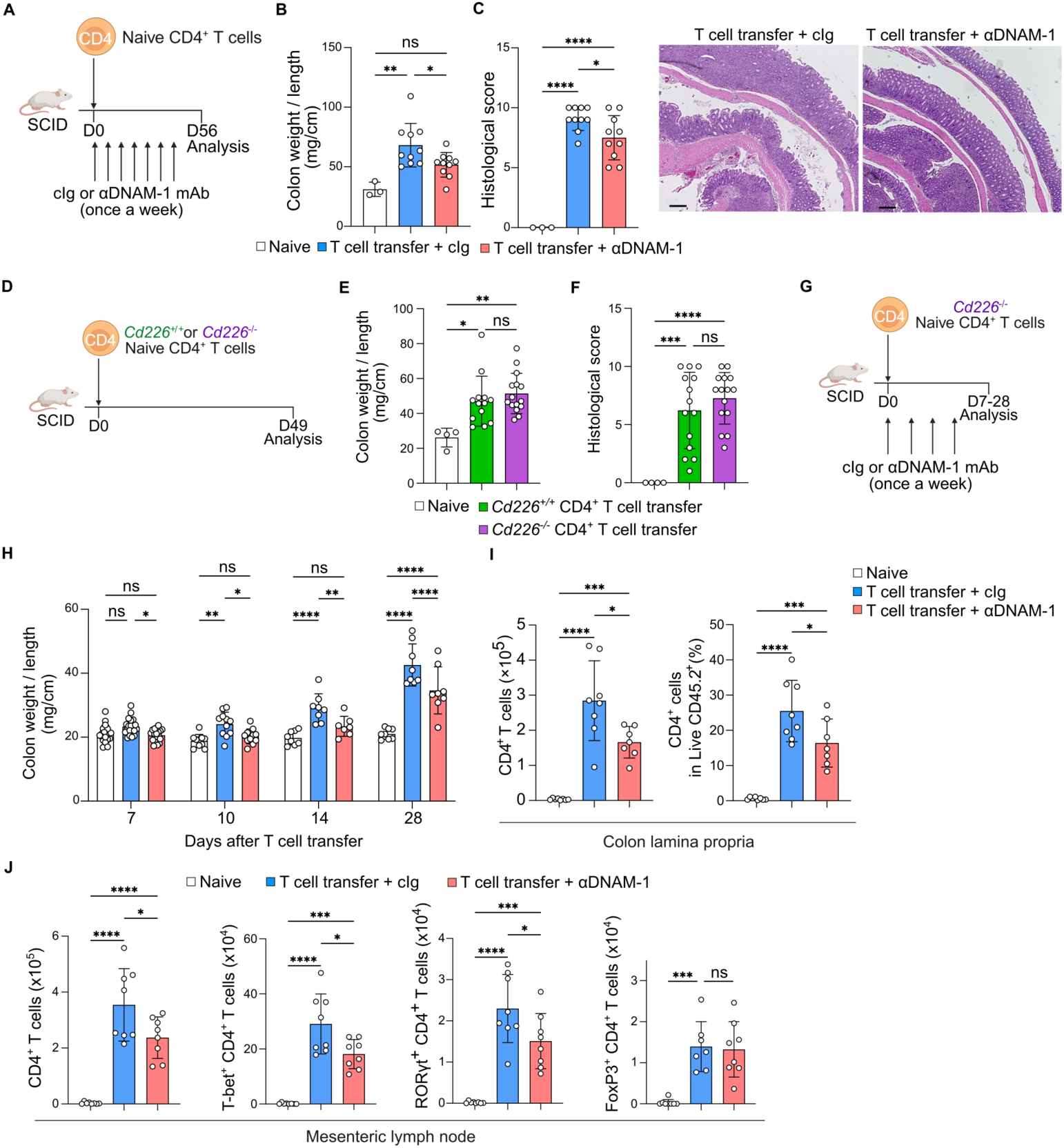
Anti–DNAM-1 mAb treatment ameliorates T cell–dependent intestinal inflammation. **(A)** Schematic illustration of the experimental design. Naive CD4⁺ T cells from BALB/c mice were adoptively transferred into SCID mice on day 0, followed by weekly intraperitoneal injections of cIg or anti–DNAM-1 mAb until one week prior to analysis on day 56. **(B)** Colon weight-to-length ratio and **(C)** histological scores and representative H&E-stained colon sections from naive mice (n = 3) and mice treated with cIg or anti–DNAM-1 mAb (n = 10) at day 56 after T cell transfer. Scale bars, 100 μm. Data were pooled from two independent experiments. **(D)** Schematic of the experimental design. Naive CD4⁺ T cells from WT or *Cd226*⁻^/^⁻ mice were adoptively transferred into SCID mice and analyzed on day 49. **(E)** Colon weight-to-length ratio and **(F)** histological scores from naive mice (n = 4) and mice transferred with WT or *Cd226*⁻^/^⁻ CD4⁺ T cells (n = 14–15) at day 49 after T cell transfer. Data were pooled from two independent experiments. **(G)** Schematic illustration of the experimental design. Naive *Cd226*⁻^/^⁻ CD4⁺ T cells were adoptively transferred into SCID mice on day 0. Recipient mice received intraperitoneal injections of cIg or anti–DNAM-1 mAb on days 0, 7, 14, and 21 (once weekly), and were analyzed at the indicated time points (days 7, 10, 14, or 28 after T cell transfer). **(H)** Colon weight-to-length ratio from naive mice (n = 8–19) and mice treated with cIg or anti–DNAM-1 mAb (n = 8–20) at days 7, 10, 14, and 28 after T cell transfer. Data at each time point were pooled from two or three independent experiments (day 7, three experiments; day 10, three experiments; day 14, two experiments; day 28, two experiments). **(I)** Absolute numbers (left) and proportion among live CD45.2⁺ cells (right) of CD4⁺ T cells in the colonic lamina propria from naive mice (n = 8) and mice treated with cIg or anti–DNAM-1 mAb (n = 8) at day 14 after T cell transfer. Data were pooled from two independent experiments. **(J)** Absolute numbers of total CD4⁺ T cells, T-bet⁺ CD4⁺ T cells, RORγt⁺ CD4⁺ T cells, and FoxP3⁺ CD4⁺ T cells in mesenteric lymph nodes from naive mice (n = 8) and mice treated with cIg or anti–DNAM-1 mAb (n = 8) at day 14 after T cell transfer. Data were pooled from two independent experiments. Data are shown as mean ± SD. Statistical significance was determined by one-way ANOVA followed by Tukey’s multiple comparisons test (B, C, E, F, I, J) or two-way ANOVA followed by Šídák’s multiple comparisons test (H); *p < 0.05, **p < 0.01, ***p < 0.001, ****p < 0.0001; ns, not significant.

To evaluate this possibility, SCID mice were reconstituted with naïve *Cd226^⁻/⁻^* CD4⁺ T cells and treated weekly with anti–DNAM-1 or control mAb (Figure 5G). Anti–DNAM-1 mAb treatment reduced colon weight-to-length ratios from days 7 to 28 (Figures 5H).

Consistently, anti–DNAM-1 mAb decreased CD4⁺ T cell infiltration in the colonic lamina propria (Figure 5I) and the total number of CD4⁺ T cells in the mesenteric lymph nodes (mLNs) at day 14 (Figure 5J). Furthermore, the absolute counts of T-bet⁺ and RORγt⁺ CD4⁺ T cells, representing Th1 and Th17 subsets, respectively, were reduced in mLNs of anti–DNAM-1 mAb–treated mice compared with controls (Figure 5J; Supplemental Figure 5E), correlating with colonic disease severity. Collectively, these results indicate that DNAM-1 expressed on innate immune cells of the recipient drives the differentiation of transferred naïve CD4⁺ T cells into pathogenic Th1 and Th17 cells, contributing to T cell–dependent intestinal inflammation.

### DNAM-1 on DCs exacerbates T cell–dependent intestinal inflammation

To assess whether ILC3s contribute to T cell–dependent colitis, we analyzed cytokine production of intestinal ILC3s 10 days after adoptive transfer of *Cd226^⁻/⁻^* CD4⁺ T cells in mice treated with control or anti–DNAM-1 mAb. Cytokine levels, including IL-22, IL-17, GM-CSF, and IFN-γ, were comparable between groups (Supplemental Figure 6A), indicating that DNAM-1 on ILC3s is dispensable in this model.

Since DNAM-1 is expressed on intestinal DCs, we examined its role in T cell–dependent colitis. DNAM-1 expression was upregulated in the CD11b⁺CD103⁻ DC subset 14 days post-T cell transfer (Figure 6A; Supplemental Figure 6B), a population critical for antigen capture and presentation ^34, 35^. Blockade of DNAM-1 reduced IL-12 production and expression of costimulatory molecules CD80, CD86, and CD40 in these DCs (Figure 6B, Supplemental Figure 6C), suggesting that DNAM-1 promotes DC activation during colitis.

**Figure 6.**
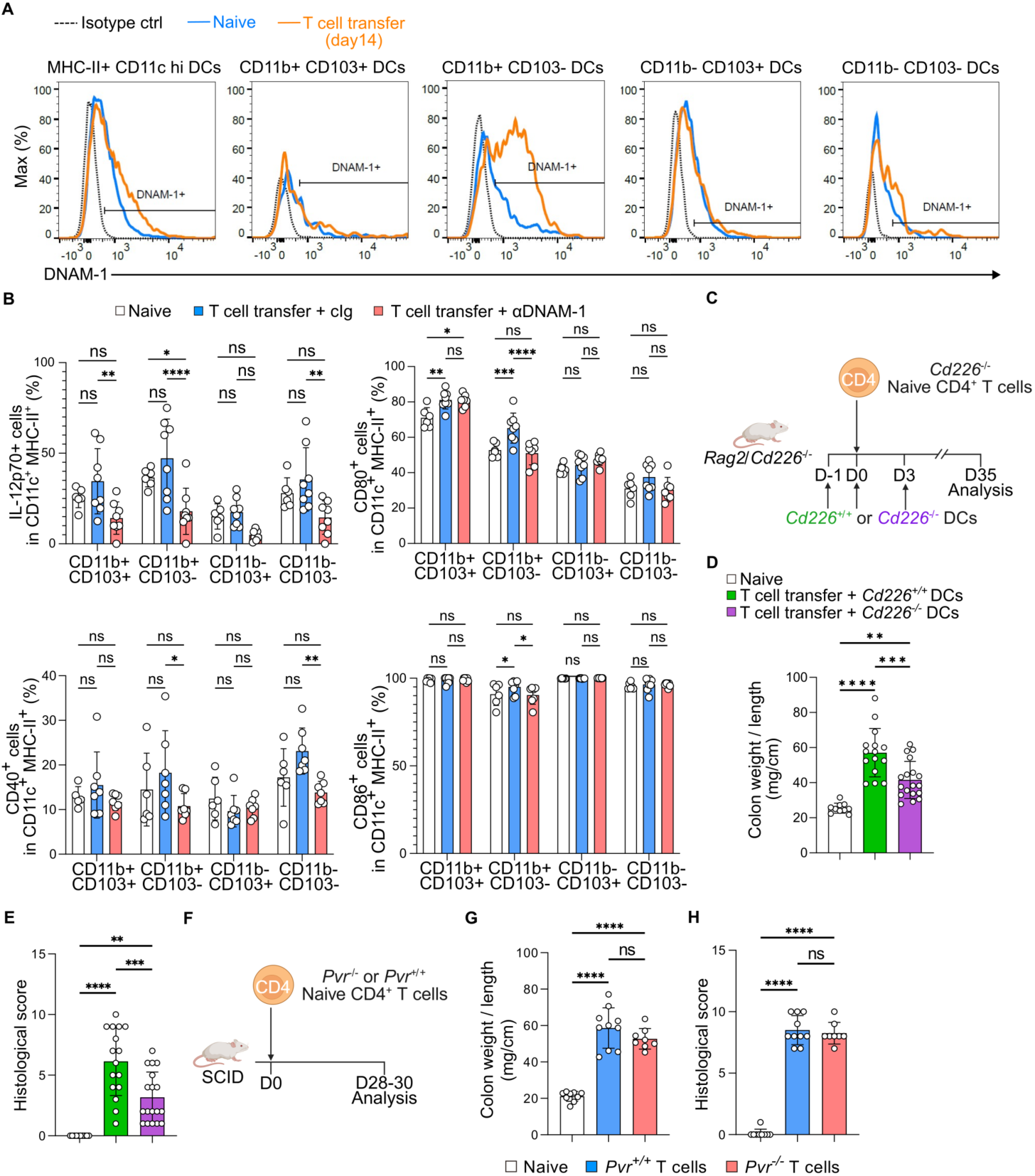
DNAM-1 on DCs exacerbates T cell–dependent intestinal inflammation. **(A)** Representative histograms of DNAM-1 expression on colonic DC subsets (MHC-II⁺ CD11cʰ^i^, further divided by CD11b and CD103 expression) from naive mice (blue) and colitic mice at day 14 after T cell transfer (orange). Isotype control staining is shown as a dashed line. **(B)** Frequencies of IL-12p70⁺, CD80⁺, CD40⁺, and CD86⁺ cells in each DC subset from the colon of naive mice (n = 6) and mice treated with cIg or anti–DNAM-1 mAb (n = 8) at day 7 after T cell transfer. Data were pooled from two independent experiments. **(C)** Schematic of the experimental design. *Cd226*⁻^/^⁻ *Rag2*⁻^/^⁻ mice received MHC-IIʰⁱ CD11cʰⁱ DCs from WT or *Cd226*⁻^/^⁻ mice on days −1, 0, and 3, and naive Cd226⁻^/^⁻ CD4⁺ T cells on day 0, followed by analysis on day 35. **(D)** Colon weight-to-length ratio and **(E)** histological scores from naive mice (n = 10) and mice reconstituted with *WT* or *Cd226*⁻^/^⁻ DCs (n = 15–18) at day 35 after T cell transfer. Data were pooled from three independent experiments. **(F)** Schematic of the experimental design. SCID mice received naive WT or *Pvr*⁻^/^⁻ CD4⁺ T cells on day 0, followed by analysis on day 28-30. **(G)** Colon weight-to-length ratio and **(H)** histological scores from naive mice (n = 9) and mice transferred with WT or *Pvr*⁻^/^⁻ CD4⁺ T cells (n = 8–10) at day 30 after T cell transfer. Data were pooled from two independent experiments. Data are shown as mean ± SD. Statistical significance was determined by one-way ANOVA followed by Tukey’s multiple comparisons test (D–G) or two-way ANOVA followed by Šídák’s multiple comparisons test (B); *p < 0.05, **p < 0.01, ***p < 0.001, ****p < 0.0001; ns, not significant.

To directly test this, *Cd226⁻^/^⁻Rag2⁻^/^⁻* mice received naïve *Cd226⁻^/^⁻* CD4⁺ T cells along with WT or *Cd226^⁻/⁻^* MHC-II^hi^ CD11c^hi^ DCs (Figure 6C). Mice receiving *Cd226⁻^/^⁻* DCs displayed reduced colon weight-to-length ratios and improved histology compared with those receiving WT DCs (Figures 6D, E), demonstrating that DNAM-1 on DCs exacerbates intestinal inflammation.

To examine whether DNAM-1 on DCs interacts with CD155 on CD4⁺ T cells and is involved in the exacerbation of intestinal inflammation, naïve WT or *Pvr⁻/⁻* CD4⁺ T cells were transferred into SCID mice. However, both groups exhibited comparable colitis (Figure 6F, G, Supplemental Figure 6D), suggesting that DNAM-1 on DCs promotes disease indirectly, likely via interactions with CD155 on intestinal cells other than transferred T cells.

Collectively, these findings indicate that DNAM-1–mediated DC activation contributes to T cell–dependent intestinal inflammation.

### Anti-DNAM-1 mAb has an additive effect with anti-TNF mAb on intestinal inflammation

Anti-TNFα mAb is among the most widely used biologics in IBD therapy. To compare the therapeutic effects of anti-DNAM-1 and anti-TNF mAbs, SCID mice were injected with anti-CD40 mAb along with control Ig, anti-DNAM-1, anti-TNF, or a combination of both antibodies. Both mAbs individually ameliorated intestinal inflammation, as evidenced by reduced colon weight-to-length ratios and histological scores (Figures 7A, Supplemental Figure 7A, B). Notably, the combination of anti-DNAM-1 and anti-TNF mAbs further lowered histological scores compared with either mAb alone (Figure 7A).

**Figure 7.**
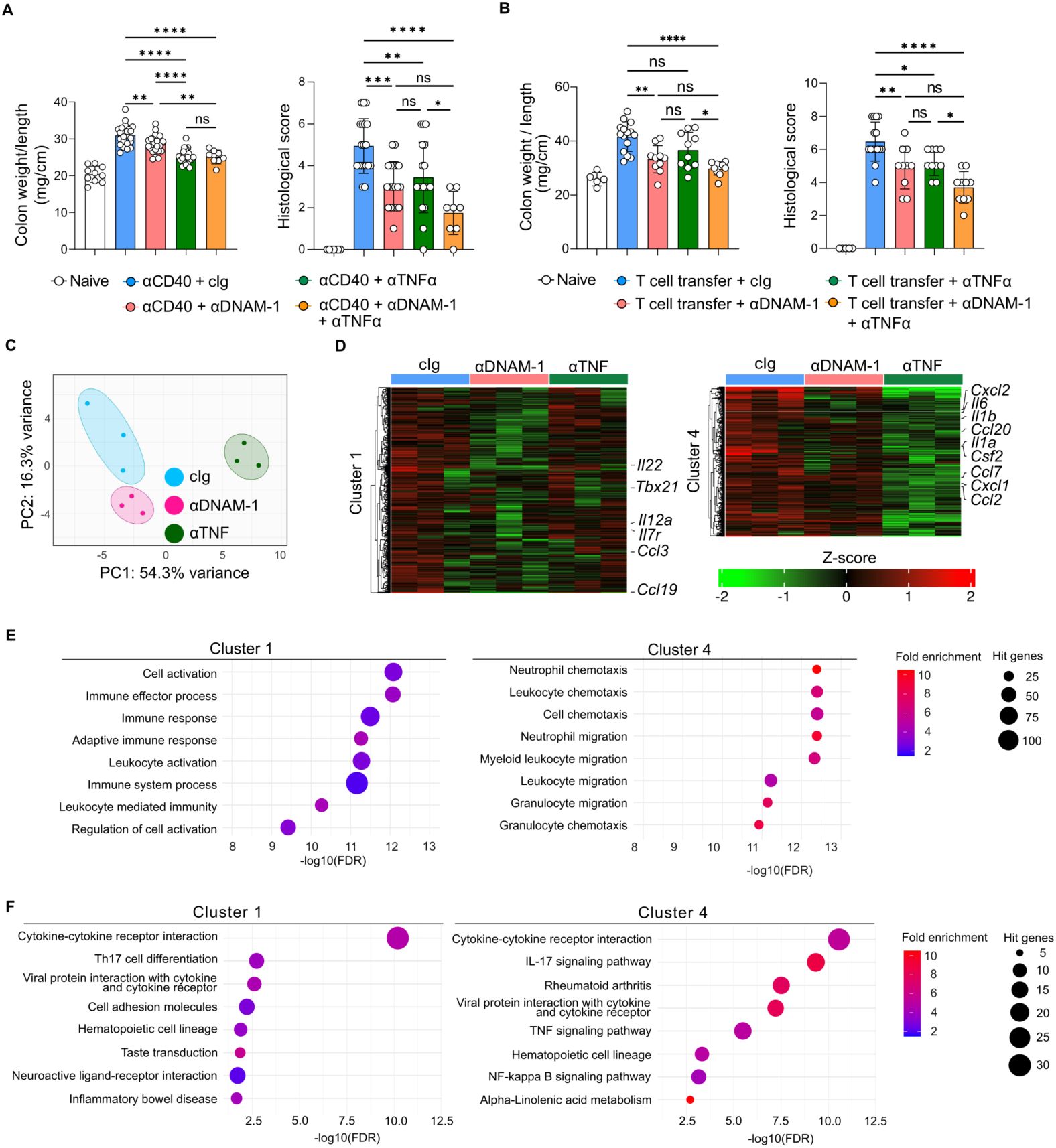
Anti-DNAM-1 mAb has an additive effect with anti-TNF mAb on intestinal inflammation. **(A)** Colon weight-to-length ratio and histological scores from naive mice (n = 10) and mice treated with cIg, anti–DNAM-1 mAb, anti-TNFα mAb, or a combination of anti–DNAM-1 and anti-TNFα mAbs (n = 8–19) at day 5 after anti-CD40 mAb injection. **(B)** Colon weight-to-length ratio and histological scores from naive mice (n = 4) and mice treated with cIg, anti–DNAM-1 mAb, anti-TNFα mAb, or a combination of anti–DNAM-1 and anti-TNFα mAbs (twice weekly; n = 10–15) at day 28 after T cell transfer. Data were pooled from three independent experiments (A, B). **(C)** Principal component analysis (PCA) of RNA sequencing data from colonic tissues of mice treated with cIg (blue), anti–DNAM-1 mAb (pink), or anti-TNFα mAb (green) at day 3 after anti-CD40 mAb injection (n = 3 per group). **(D)** Heatmap of differentially expressed genes identified by RNA sequencing of colonic tissues from mice treated with cIg, anti–DNAM-1 mAb, or anti-TNFα mAb (n = 3 per group) at day 3 after anti-CD40 mAb injection. Genes were grouped by hierarchical clustering, and representative genes in Cluster 1 and Cluster 4 are indicated. Color scale represents Z-score of normalized expression values. **(E)** Gene ontology (GO) biological process enrichment analysis of genes in Cluster 1 (left) and Cluster 4 (right). Dot size represents the number of hit genes; color represents fold enrichment. **(F)** KEGG pathway enrichment analysis of genes in Cluster 1 (left) and Cluster 4 (right). Dot size represents the number of hit genes; color represents fold enrichment. Data in (A, B) are shown as mean ± SD. Statistical significance was determined by one-way ANOVA followed by Tukey’s multiple comparisons test (A–D); *p < 0.05, **p < 0.01, ***p < 0.001, ****p < 0.0001; ns, not significant.

Similarly, in the T cell–dependent colitis model induced by adoptive transfer of naïve CD4⁺ T cells, administration of either anti-DNAM-1 or anti-TNF mAb twice weekly reduced colon weight-to-length ratios and histological scores at day 28, and the combination of both antibodies produced an additive effect (Figure 7B, Supplemental Figures 7C, D). These findings suggest that anti-DNAM-1 mAb exerts therapeutic effects through a mechanism distinct from anti-TNF mAb.

To investigate the molecular basis for this distinction, RNA sequencing was performed on inflamed colonic tissues from mice with anti-CD40 mAb-induced colitis treated with each antibody. Principal component analysis revealed clear separation between anti-DNAM-1 and anti-TNF treatment groups (Figure 7C), and clustering analysis identified five gene expression patterns (Figure 7D, Supplemental Figure 7E). Cluster 1 genes were specifically downregulated in the anti-DNAM-1 group, whereas cluster 4 genes were downregulated in the anti-TNF group. Gene ontology analysis indicated that cluster 1 genes were enriched for “cell activation” and “immune effector process,” while cluster 4 genes were associated with “neutrophil chemotaxis” and “leukocyte migration” (Figure 7E). KEGG pathway analysis further highlighted that anti-DNAM-1 preferentially modulated pathways such as Th17 cell differentiation, cell adhesion molecules, and IBD, whereas anti-TNF predominantly affected NF-κB and IL-17 signaling pathways (Figure 7F). In contrast, the other cluster genes were not found to be associated with the pathology of intestinal inflammation (Supplemental Figures 7F, G). Collectively, these results indicate that anti-DNAM-1 and anti-TNF antibodies regulate intestinal inflammation via complementary mechanisms, supporting the potential of anti-DNAM-1 as a novel therapeutic strategy targeting inflammatory programs not fully addressed by anti-TNF therapy.

## Discussion

Our study identifies DNAM-1 (CD226) as a central regulator of intestinal inflammation that operates through distinct cell-type-specific mechanisms in innate and adaptive immune contexts. We demonstrate that DNAM-1 promotes ILC3-mediated inflammation in an innate immune–driven colitis model by amplifying IL-22 and GM-CSF production via the Akt–mTORC1–HIF-1α signaling axis. Conversely, in T cell–dependent colitis, DNAM-1 expressed on dendritic cells drives pathogenic Th1 and Th17 differentiation. Importantly, blockade of DNAM-1 ameliorates disease in both settings and exhibits additive effects with anti-TNF therapy, highlighting DNAM-1 as a multifunctional immunoregulatory node bridging disparate inflammatory programs in intestinal inflammation.

A key finding of this study is the identification of DNAM-1 as a cell-intrinsic amplifier of ILC3 effector function. Although DNAM-1 has been well characterized in cytotoxic lymphocytes ^20, 36, 37^, its role in ILC3 biology has remained largely unexplored. We show that DNAM-1 is highly expressed on intestinal ILC3 subsets and quantitatively enhances IL-22 and GM-CSF production in response to IL-23. Adoptive transfer experiments in lymphocyte-deficient mice establish that DNAM-1 expression intrinsic to ILC3s is sufficient to exacerbate colitis, providing direct evidence that DNAM-1 functions as a pathogenic driver in innate immune–mediated intestinal inflammation.

Mechanistically, our data reveal that DNAM-1 engages CD155 expressed on myeloid cells to deliver a costimulatory signal that cooperates with IL-23 to selectively augment IL-22 production. This interaction activates the Akt–mTORC1–HIF-1α pathway, which has been implicated in the metabolic and transcriptional regulation of ILC3 effector programs ^27, 38, 39, 40^. Notably, DNAM-1 stimulation alone is insufficient to induce cytokine production, underscoring its role as a context-dependent amplifier rather than a primary activation signal. These findings provide a mechanistic framework explaining how IL-22, a cytokine with homeostatic functions, can be diverted into a pathogenic mediator under inflammatory conditions ^16, 41^.

In contrast to innate immune–driven colitis, DNAM-1 plays a distinct role in T cell–dependent disease. We show that DNAM-1 expression on dendritic cells, but not on CD4⁺ T cells or ILC3s, promotes colitis by enhancing IL-12 production and upregulating the costimulatory landscape (CD80, CD86, and CD40), thereby facilitating Th1 and Th17 differentiation ^42^. This unexpected requirement for DC-intrinsic DNAM-1, rather than T cell-intrinsic signaling, highlights the context- and cell type–specific functions of DNAM-1 in the gut and underscores its capacity to coordinate innate and adaptive immune responses through different cellular effectors.

The therapeutic implications of our findings are notable. DNAM-1 blockade attenuates intestinal inflammation in both innate and adaptive models of colitis and demonstrates additive effects when combined with anti-TNF therapy. Transcriptomic analyses reveal that DNAM-1 and TNF inhibition modulate largely non-overlapping inflammatory programs: TNF inhibition primarily targets NF-κB-mediated acute myeloid responses, whereas DNAM-1 blockade preferentially targets lymphocyte activation, Th17 differentiation, and cell adhesion pathways. These data suggest that targeting DNAM-1 may complement existing biologics and provide therapeutic benefit in patient populations that do not achieve sustained remission with current therapies ^43, 44^.

Several limitations warrant consideration. Our study primarily relied on antibody-mediated DNAM-1 blockade, and the long-term consequences of sustained DNAM-1 inhibition on host defense and tissue homeostasis remain to be determined ^36, 37^. In addition, while CD155 on myeloid cells appears to provide the dominant activating signal for DNAM-1, the relative contribution of epithelial versus hematopoietic ligand expression requires further investigation. Moreover, although the CD11b+CD103- DC population upregulates the expression of DNAM-1, costimulatory molecules, and IL-12 after T cell transfer, whether and how DNAM-1 on this DC population contributes to colitis remains undetermined.

Finally, validation of DNAM-1 expression and function in human IBD tissues will be essential to support clinical translation. In summary, we identify DNAM-1 as a context-dependent regulator of intestinal inflammation that bridges innate and adaptive immune programs. By amplifying ILC3-mediated cytokine production and promoting dendritic cell–driven T cell pathogenicity, DNAM-1 contributes to colitis pathogenesis. Our transcriptomic finding that *CD226* is an IL-23-responsive gene within IBD risk loci reinforces the clinical relevance of our murine models. Therapeutic targeting of DNAM-1 ameliorates disease and synergizes with anti-TNF therapy, highlighting DNAM-1 as a promising target for future IBD treatment strategies.

## Acknowledgments

This work was supported by JSPS KAKENHI (21H04836 to A.S., 21K19469 and 16H05169 to K.S, and 21K15462 to Kazuki S) and JST SPRING (JPMJSP2124 to N.I). The authors thank M. Kaneko for administrative support.

## Author Contributions

N.I. Kazuki S, K.O, M.S.A, F. A. and T-H. K. performed experiments. N.I., C. N-O, Kazuko S, and A.S. performed data analysis and project planning. N.I., M.S.A, Kazuko S., and A.S. contributed to writing the manuscript. Kazuko S and A.S. supervised the overall project.

## Declaration of Interests

F.A. is an employee and A.S. is a board member of TNAX biopharma corporation. C. N-O, Kazuko S. and A.S. own a stake in TNAX biopharma corporation.

## Materials and Methods

### Mice

SCID (NOD.CB17-*Prkdcscid*/Jcl) mice were purchased from Clea Japan (Tokyo, Japan). BALB/c mice deficient in *Rag2* (*Rag2*-/-) or both *Rag2* and IL-2 receptor common g chain (*Rag2*-/-*Il2γc*-/-) ^45^ were purchased from Jackson Laboratory. BALB/c mice deficient in DNAM-1 (*Cd226*-/-) or CD155 (*Pvr*-/-) were generated, as described previously ^37, 46^. *Rag2*-/-*Cd226*-/- BALB/c mice were generated by crossing *Rag2*-/- mice with *Cd226*-/- mice. Male and female mice aged 8-12 weeks were used for the experiments. All mice were housed and maintained under specific pathogen-free conditions. All animal experiments were conducted in accordance with the guidelines of the animal ethics committee of the Laboratory Animal Resource Center, University of Tsukuba (approval no., 22-155 and 23-120).

### Antibodies

Anti-DNAM-1 (TX42.2; mouse IgG1 containing mutated amino acids at residues 234 and 235 from L to A in the Fc portion to avoid Fcγ receptor binding) was generated by our laboratory. Anti-KLH (IC17; mouse IgG1) antibody was generated in our laboratory. Anti-TNF-α mAb (XT3.11), GM-CSF (MP1-22E9), CD40 (FGK4.5/FGK45) and Rat IgG2a (2A3), was purchased from BioXcell (Lebanon, NH). Anti-TNF-α mAb (XT3.11) was purchased from Selleck (Houston, USA). mAbs against mouse CD49b (DX5), CD11b (M1/70), CD86 (GL1), CD155 (3F1), rat IgG2a, rat IgG1, hamster IgG1, hamster IgG2 (KKLH), rat IgG2a (RTK2758), Ly-6G (1A8), CD40 (3/23), CD112 (829038), Gr-1 (RB6-8C5), CD90.2 (53-2.1), CD11c (HL3), CD11b (M1/70), Ly-6C (AL-21), NKp46 (29A1.4), RORγt (Q31-378), IFN-γ (XMG1.2), IL-17A (TC11-18H10), IL-12(p40/p70) (C11.5), human CD19 (HIB19), CD138 (MI15), CD117 (104D2) and Fluorochrome-conjugated streptavidin were purchased from BD Biosciences (Franklin Lakes, NJ). mAb against T1/ST2 (DJ8) was purchased from MD Bioproducts (St. Paul, MN). mAb against mouse IL-22 (1H8PWSR), CD25 (7D4),T-bet (4B10) GATA-3 (TWAL), HIF-1 alpha (Mgc3), and IL-22 (IL22JOP), and human CD127 (eBioRDR5) was purchased from Invitrogen (Waltham, MA). mAbs against mouse TCR-β (H57-597), CD11b (M1/70), CD155 (TX56), KLRG1 (2F1), CD25 (PC61), NKp46 (29A1.4), Ly-6G (IA8), Gr-1 (RB6-8C5), CD80 (B7-1), CD103 (2E7), CD49b (DX5), CD45RB (C363-16A), CD90.2 (53-2.1), I-A/I-E (M5/114.15.2), CD45.2 (104), CD45 (30-F11), CD64 (X54-5/7.1), CD170/Siglec-F (S17007L), CD11c (N418), CD4 (RMA-5), CD4 (GK1.5), F4/80 (BM8), CD127(A7R34), CD11b (M1/70), Ly-6C (HK1.4), CCR6 (29-2L17), T-bet (4B10), IFN-γ (XMG1.2), GM-CSF (MP1-22E9), human CD161 (HP-3G10), CCR6 (G034E3), NKp44 (P44-8), CRTH2 (BM16), and Fluorochrome-conjugated streptavidin were purchased from Biolegend (San Diego, CA).

mAb against phospho-Akt(T308) (D25E6) and phospho-S6 ribosomal protein (s235/236) (D57.2.2E) were purchased from Cell Signaling Technology (Massachusetts, USA). mAb against human CD3 was purchased from Cytec Biosciences (California, USA). mAb against human CD16 and FcεRIα were purchased from Miltenyi biotec.

### Isolation of Lamina Propria Lymphocytes (LPLs) and ILC3s

Small intestine and colon tissues were processed to isolate lamina propria leukocytes (LPLs). Tissues were washed with cold PBS, cut longitudinally, and incubated in HBSS containing 5 mM EDTA and 1 mM DTT to remove epithelial cells. Remaining tissues were digested in RPMI-1640 with 1 mg/mL collagenase D (Roche) and 0.1 mg/mL DNase I (Sigma-Aldrich) at 37°C for 30 min. Cells were suspended in 40% Percoll PLUS (GE Healthcare), overlaid onto 80% Percoll PLUS, and centrifuged at 860 g for 30 min at 21°C. The buffy coat layer was collected, washed with PBS containing 2 mM EDTA, followed by RPMI-1640, and used as LPLs. LPLs were stained with the following antibodies: CD45.2, CD11b, CD11c, KLRG1, CD49b, CD90.2, and CD127. ILC3s were identified as PI-CD45.2^+^ Lin^-^ (CD11b, CD11c, KLRG1, CD49b) CD90.2^+^ CD127^+^ and sort-purified using FACSAria III or FACS Aria SORP (BD). Purity exceeded 95%.

### Human peripheral blood ILC3 analysis

Peripheral blood samples (50 mL) were obtained from two healthy volunteers and collected in heparinized tubes. Whole blood was diluted at 1:1 with PBS, carefully layered onto Lymphoprep (STEMCELL Technologies, Canada Vancouver), and centrifuged at 800 × g for 20 min at room temperature without brake or acceleration. The buffy coat was collected and washed with PBS. Cells were then incubated with anti-human CD138-FITC, anti-human FcεRIα-FITC, and anti-human CD3-FITC for 30 min on ice. After washing, cells were incubated with CD11b, CD34, CD56, CD14, CD16, and CD19 microbeads together with anti-FITC microbeads (Miltenyi) for 15 min at 4°C, and unwanted cell populations were depleted using MACS.

The enriched cell fraction was incubated with human FcR Blocking Reagent (Miltenyi) for 10 min on ice, followed by staining with the antibody cocktail for 60 min on ice. After washing with PBS, fluorochrome-conjugated streptavidin was added and incubated for 30 min on ice. Cells were then washed and analyzed by flow cytometry.

Human ILC3s were defined as Lin^-^ (CD3, CD11b, CD14, CD16, CD19, CD138, and FcεRIα) CD127^+^CD161^+^CD117^+^CRTH2^-^ cells. ILC3 subsets were further classified as NKp44^+^CCR6^+^, NKp44^+^CCR6^-^, and NKp44^-^CCR6^-^ populations.

### In Vitro Stimulation of ILC3s

Sorted ILC3s (3–8 x 10^3^ cells) were stimulated in 96-well plates coated with 1 mg/mL DOTAP (Cayman Chemical). Anti-DNAM-1 mAb (clone TX42.2, 1 μg) or isotype control IgG (clone IC17, 1 μg) was added in 50 mM carbonate buffer (pH 9.6) and incubated overnight at 4°C. Plates were washed with PBS and blocked with complete RPMI-1640 for 30 min at room temperature. ILC3s were cultured in the presence or absence of rmIL-23 (1 ng/mL) and inhibitors including Akt inhibitor (100 or 1 μM, Abcam), Rapamycin (10 or 1 ng/mL, Sigma-Aldrich), or HIF-1α inhibitor (FM19G11, 1 or 0.1 μM, Selleck) in complete RPMI-1640 supplemented with 10% FBS, 50 μM 2-mercaptoethanol, 2 mM L-glutamine, 100 U/mL penicillin, 0.1 mg/mL streptomycin, 10 mM HEPES, 1 mM sodium pyruvate, and 100 μM MEM non-essential amino acids under 5% CO2 at 37°C. Culture supernatants were collected for ELISA (Thermo Fisher Scientific) or cytometric bead array (CBA) (BD Biosciences), and cells were processed for qRT-PCR or Phosflow analysis of p-Akt, p-S6, and HIF-1α.

### Isolation of Myeloid Cells and ILC3–Myeloid Co-culture

Myeloid cells were isolated from naïve or inflamed intestines 3 days post anti-CD40 mAb treatment by positive selection with CD11b or CD11c microbeads and LS columns (Miltenyi Biotec). ILC3s were sorted from the flow-through fraction. WT or *Cd226*^-/-^ ILC3s (1 x 10^4^) and WT, *Cd226*^-/-^, or *Pvr^-^*^/-^ myeloid cells (2 x 10^4^) were co-cultured in 96-well plates for 24 h in complete RPMI-1640 medium in the presence or absence of IL-23 (1 ng/mL) and anti-DNAM-1 mAb (10 μg/mL) or isotype control IgG (10 μg/mL). Supernatants were collected for ELISA or CBA, and cells were dissolved in ISOGEN (Nippon Gene) for qRT-PCR.

### ELISA and Cytometric Bead Array (CBA)

Mouse IL-22 in culture supernatants was quantified using a Mouse IL-22 Uncoated ELISA Kit (Thermo Fisher Scientific) according to the manufacturer’s protocol. IFN-γ, GM-CSF, IL-17A, TNF, IL-1β, and IL-6 were measured by CBA (BD Biosciences).

### Adoptive Transfer of ILC3s

WT or *Cd226*^-/-^ ILC3s (2 x 10^5^) from the small intestine or colon were adoptively transferred via intravenous injection into 8-week-old *Rag*2^-/-^*γc*^-/-^ BALB/c mice. On day 14 post-transfer, 200 μg anti-CD40 mAb was administered intraperitoneally to induce colitis.

### Histopathology and Ex Vivo Colon Culture

Colons were longitudinally opened, rolled into “Swiss rolls,” fixed in formalin, embedded in paraffin, and sectioned at 4–5 μm. Sections were stained with hematoxylin and eosin (H&E), and images were acquired using a BZ-X710 microscope (Keyence).

Histopathological scoring of epithelial damage and inflammatory cell infiltration was performed as described ^47^. For ex vivo culture, colon tissues were flushed with cold PBS, cut into pieces, and incubated in 1 mL RPMI-1640 supplemented with 10% FBS, 100 IU/mL penicillin, 100 μg/mL streptomycin, 1 mM sodium pyruvate, and 2 mM L-glutamine in 24-well plates for 16–20 h at 37°C under 5% CO2. Supernatants were collected for CBA.

### Quantitative Real-Time PCR (qRT-PCR)

Total RNA was extracted from colon tissues, FFPE sections, or cells using ISOGEN (Nippon Gene) or the RecoverAll™ Total Nucleic Acid Isolation Kit (Thermo Fisher Scientific) for FFPE samples. cDNA was synthesized using the High-Capacity cDNA Reverse Transcription Kit (Thermo Fisher Scientific). qRT-PCR was performed on the ABI 7500 Fast Real-Time PCR System using Power SYBR Green PCR Master Mix (Thermo Fisher Scientific) and 50 nM primers. Gene expression was normalized to *Actb*. PCR primers used were as follows: Csf2-F, 5’-CCTGGGCATTGTGGTCTACAG-3’; Csf2-R, 5’-GGTTCAGGGCTTCTTTGATGG-3’; Infg-F, 5’-ACAGCAAGGCGAAAAAGGATG-3’; Infg-R, 5’-TGGTGGACCACTCGGATGA-3’; Il1b-F, 5’-GAAATGCCACCTTTTGACAGTG-3’; Il1b-R, 5’-TGGATGCTCTCATCAGGACAG-3’; Il6-F, 5’-GAGGATACCACTCCCAACAGACC-3’; Il6-R, 5’-AAGTGCATCATCGTTGTTCATACA-3’; Il12p35-F, 5’-CCCTTGCCCTCCTAAACCAC-3’; Il12p35-R, 5’-AAGGAACCCTTAGAGTGCTTACT-3’; Il12p40-F, 5’-CAGAAGCTAACCATCTCCTGGTTTG-3’; Il12p40-R, 5’-CCGGAGTAATTTGGTGCTCCACAC-3’; Il13-F, 5’-TGAGGAGCTGAGCAACATCAC-3’; Il13-R, 5’-GGTTACAGAGGCCATGCAAT-3’; Il22-F, 5’-TTGAGGTGTCCAACTTCCAGCA-3’; Il22-R, 5’-AGCCGGACGTCTGTGTTGTTA-3’; Il23a-p19-F, 5’-TGCTGGATTGCAGAGCAGTTAA-3’; Il23a-p19-R, 5’-GCATGCAGAATTCCGAAGA-3’; Rorc-F, 5’-GGAGGACAGGGAGCCAAGTT-3’; Rorc-R, 5’-CCGTAGTGGATCCCAGATGACT-3’; Tbx21-F, 5’-AGCAAGGACGGCGAATGTT-3’; Tbx21-R, 5’-GGGTGGACATATAAGCGGTTC-3’; Tnfa-F, 5’-GGGCCACCACGCTCTTC-3’; Tnfa-R, 5’-GGTCTGGGCCATAGAACTGATG-3’; Cxcl1-F, 5’-CCGAAGTCATAGCCACACTCAA-3’; Cxcl1-R, 5’-GCAGTCTGTCTTCTTTCTCCGTTA-3’; Cxcl2-F, 5’-GAAGTCATAGCCACTCTCAAGG-3’; Cxcl2-R, 5’-CCTCCTTTCCAGGTCAGTTAGC-3’; Cxcl16-F, 5’-TTATCAGGTTCCAGTTGCAGTC-3’; Cxcl16-R, 5’-TGGTGGTGAAAACTCTTCCCA-3’; Ccl2-F, 5’-GCTGGAGCATCCACGTGTT-3’; Ccl2-R, 5’- TTGGGATCATCTTGCTGGTGA-3’; Actb-F, 5’-ACTGTCGAGTCGCGTCCA-3’; Actb-R, 5’-GCAGCGATATCGTCATCCAT-3’.

### Bulk RNA Sequencing

Total RNA from colonic tissues of anti-CD40 mAb-induced colitic mice treated with control, anti-DNAM-1, or anti-TNFα antibodies was processed by Novogene. mRNA was enriched using poly-T oligo magnetic beads, fragmented, and converted into directional cDNA libraries. Libraries were sequenced on the MGI DNBSEQ-T7 platform (paired-end 150 bp, ∼20 million reads/sample). Raw reads were quality filtered, aligned to mm10 using HISAT2^48^, and counted with FeatureCounts^49^. Genes with <10 total counts were excluded. Differential expression analysis and normalization were performed using DESeq2^50^ in R (v4.4.1)^51^. PCA plots were generated using ggplot2^52^ and ggforce.

### Single-Cell RNA Sequencing Analysis of ILC3s

IL-23-stimulated and unstimulated mouse ILC3 scRNA-seq data (GEO: GSE229976) were analyzed in Seurat (v5.2.1)^53–57^. Cells with 1,000–5,000 genes and <10% mitochondrial content were retained. Data were normalized, integrated, clustered, and projected using PCA and UMAP. Non-ILC3 clusters (*Cd3e*^+^, *Mki67*^+^, or *Il23r*^-^) were excluded. Differentially expressed genes (adjusted p < 0.01, |log2FC| > 1) were identified using the Wilcoxon rank-sum test.

### Immunohistochemistry (IHC)

Neutrophils and monocytes were stained with Ly-6G (clone E6Z1T; Cell Signaling Technology) and Ly-6C (clone RM1151; Abcam), respectively. Images were acquired randomly with the Mantra 2™ Quantitative Pathology Workstation (PerkinElmer, x40). Cells were manually counted in a blinded manner.

### Statistical Analysis

Data are presented as mean ± SD. Comparisons were made using one-way ANOVA with post hoc testing or two-tailed Student’s t-test (GraphPad Prism 10). P-values < 0.05 were considered statistically significant. Statistical outliers were excluded using the interquartile range (IQR) method.

