## Supplemental figure for "DNAM-1 immunoreceptor integrates innate and adaptive immune programs to drive intestinal inflammation"

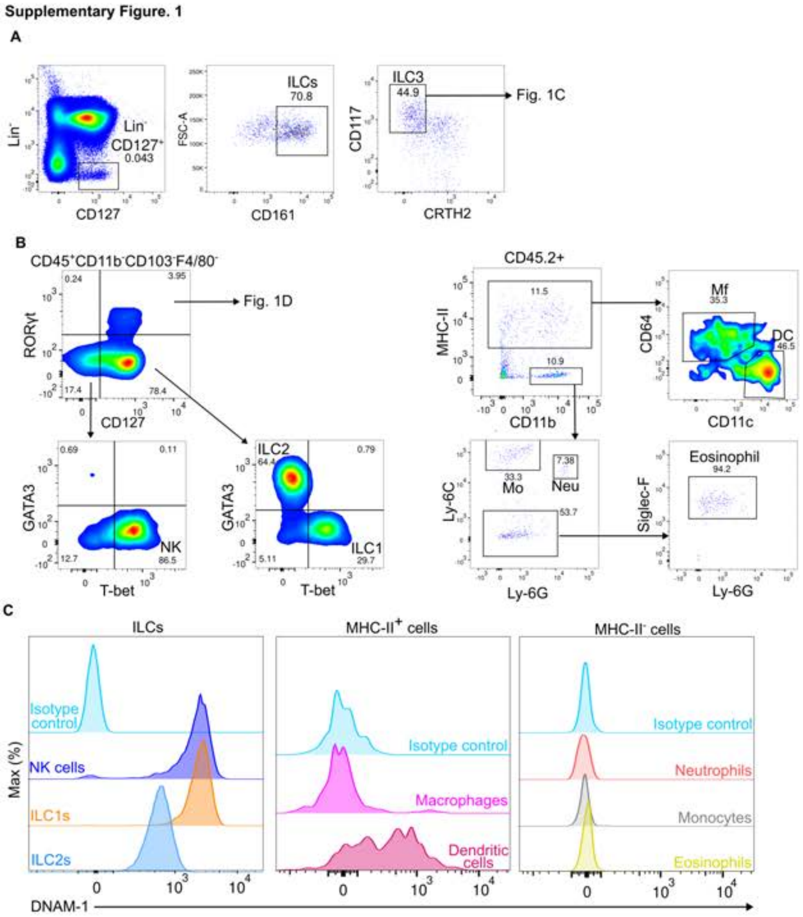


### Supplementary Figure 1.

**(A)** Gating strategy for human ILC3s in the peripheral blood. Lin$^-$ CD127$^+$ CD161$^+$CD117$^+$ CRTH2$^-$ cells were designated as ILC3c (see Figure 1C). **(B)** Gating strategy for mouse colon ILC subsets and myeloid cell populations. CD45$^+$ CD11b$^-$ CD103$^-$ F4/80$^-$ cells were gated for RORγt$^+$ CD127$^+$ ILC3s (see Figure 1D), and further subdivided by GATA3 and T-bet expression to identify NK cells (CD127$^-$ T-bet$^+$ GATA3$^-$), ILC1s (CD127$^+$ T-bet$^+$ GATA3$^-$ RORγt$^-$), and ILC2s (CD127$^+$ T-bet$^-$ RORγt$^-$ GATA3$^+$). CD45.2$^+$ cells were gated by MHC-II and CD11b expression, then further classified into macrophages (Mf; CD64$^+$ CD11c$^-$), dendritic cells (DC; CD64$^-$ CD11c$^{hi}$), monocytes (Mo) and neutrophils (Neu) by Ly-6C and Ly-6G expression, and eosinophils (Siglec-F$^+$). **(C)** Representative histograms of DNAM-1 expression on mouse ILCs (NK cells, ILC1s, ILC2s), MHC-II$^+$ cells (macrophages, dendritic cells), and MHC-II$^{-}$ cells (neutrophils, monocytes, eosinophils) from the colon of naive SCID mice. Isotype control staining is shown.

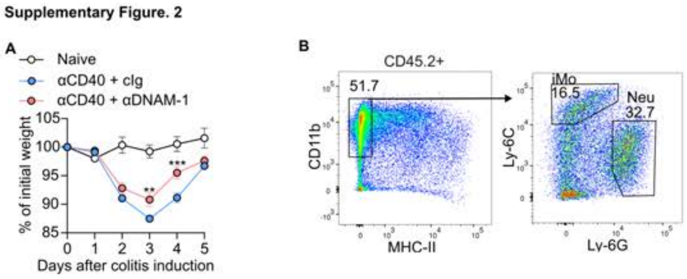


**Supplementary Figure 2.**

**(A)** Body weight changes (% of initial weight) in naive mice (n = 5) and mice treated with cIg or anti–DNAM-1 mAb (n = 14) following anti-CD40 mAb injection. Data were pooled from two independent experiments. Data are shown as mean ± SD. Statistical significance was determined by two-way ANOVA followed by Šídák's multiple comparisons test; ***$p < 0.001$. **(B)** Gating strategy for infiltrating neutrophils (Neu; MHC-II$^-$ CD11b$^+$ Ly-6G$^+$) and inflammatory monocytes (iMo; MHC-II$^-$ CD11b$^+$ Ly-6C$^+$) within CD45.2$^+$ colonic cells (see Figure 2E).

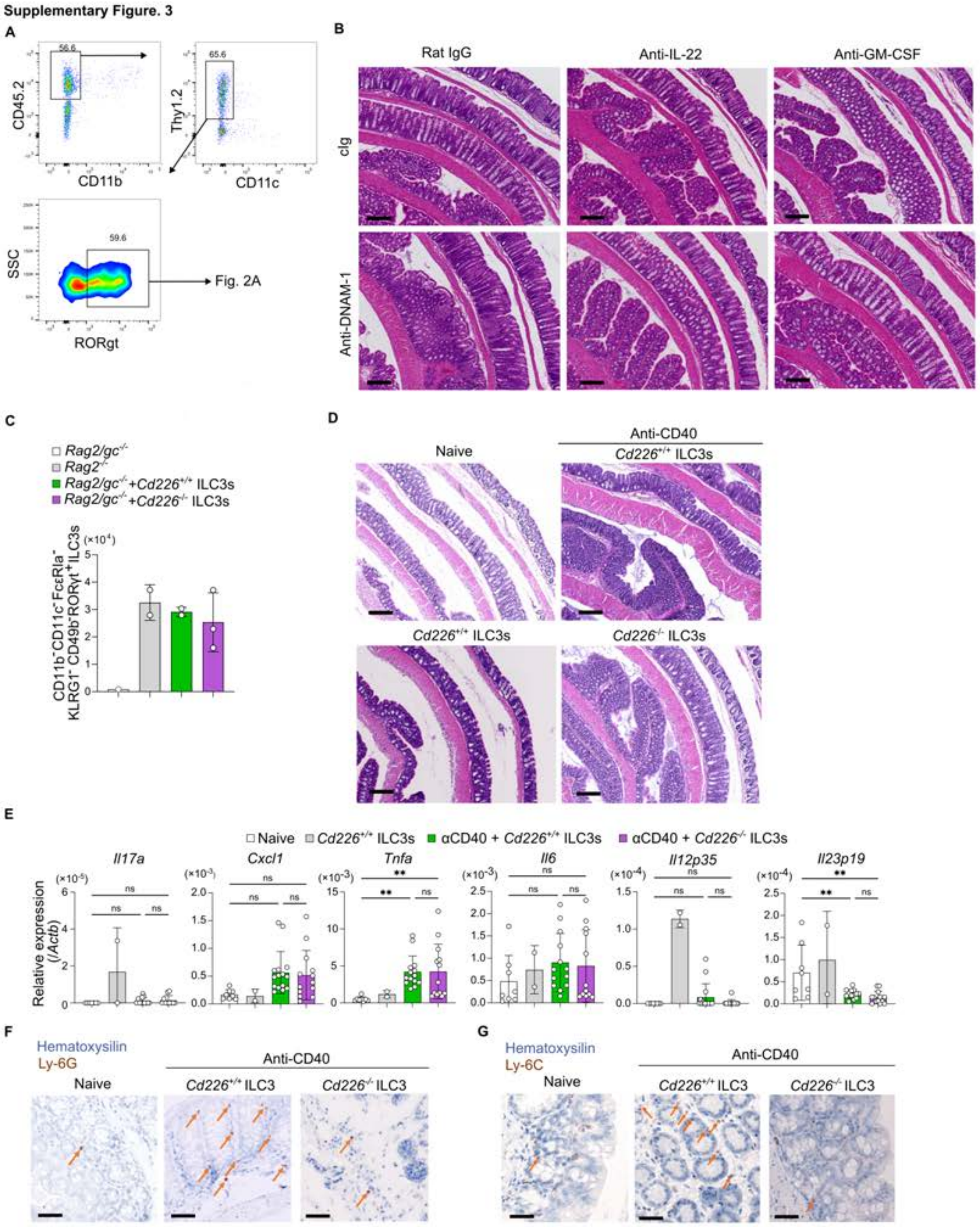


**Supplementary Figure 3.**

**(A)** Gating strategy for intestinal ILC3s used in Figure 3A. CD45.2$^+$ CD11b$^-$ CD11c$^-$ Thy1.2$^+$ cells were gated, and RORγt$^+$ cells were identified as ILC3s. **(B)** Representative H&E-stained colon sections from mice co-treated with anti-CD40 mAb, either cIg or anti-DNAM-1 mAb, together with rat IgG, anti–IL-22 mAb, or anti–GM-CSF mAb (see Figures 3B, C). Scale bars, 100 μm. **(C)** Absolute numbers of colonic ILC3s (CD11b$^-$ CD11c$^-$

FcεRIα$^{-}$ KLRG1$^{-}$ CD49b$^{-}$ RORγt$^{+}$) in *Rag2*$^{-/-}$*γc*$^{-/-}$ mice (n = 1), *Rag2*$^{-/-}$ mice (n = 2), and Rag2$^{-/-}$γc$^{-/-}$ mice reconstituted with WT (n = 2) or *Cd226*$^{-/-}$ (n = 3) ILC3s at 14 days after ILC3 transfer without colitis induction. Data are representative of two independent experiments. **(D)** Representative H&E-stained colon sections from naive mice and mice reconstituted with WT or *Cd226*$^{-/-}$ ILC3s at day 5 after anti-CD40 mAb injection (see Figures 3E, F). Scale bars, 100 μm. **(E)** Relative mRNA expression of indicated genes (normalized to Actb) in the colon from naive mice (n = 8), mice reconstituted with WT ILC3s without colitis induction (n = 2), and mice reconstituted with WT (n = 15) or *Cd226*$^{-/-}$ (n = 13) ILC3s at day 5 after anti-CD40 mAb injection. RNA was extracted from FFPE colon sections. Data were pooled from four independent experiments. **(F, G)** Representative immunohistochemical staining for (F) Ly-6G (neutrophils) and (G) Ly-6C (inflammatory monocytes) in colon sections from naive mice and mice reconstituted with WT or *Cd226*$^{-/-}$ ILC3s at day 5 after anti-CD40 mAb injection (see Figure 3H). Arrows indicate positively stained cells. Scale bars, 40 μm. Data in (C, E) are shown as mean ± SD. Statistical significance in (E) was determined by one-way ANOVA followed by Tukey's multiple comparisons test; $^{**}p < 0.01$; ns, not significant.

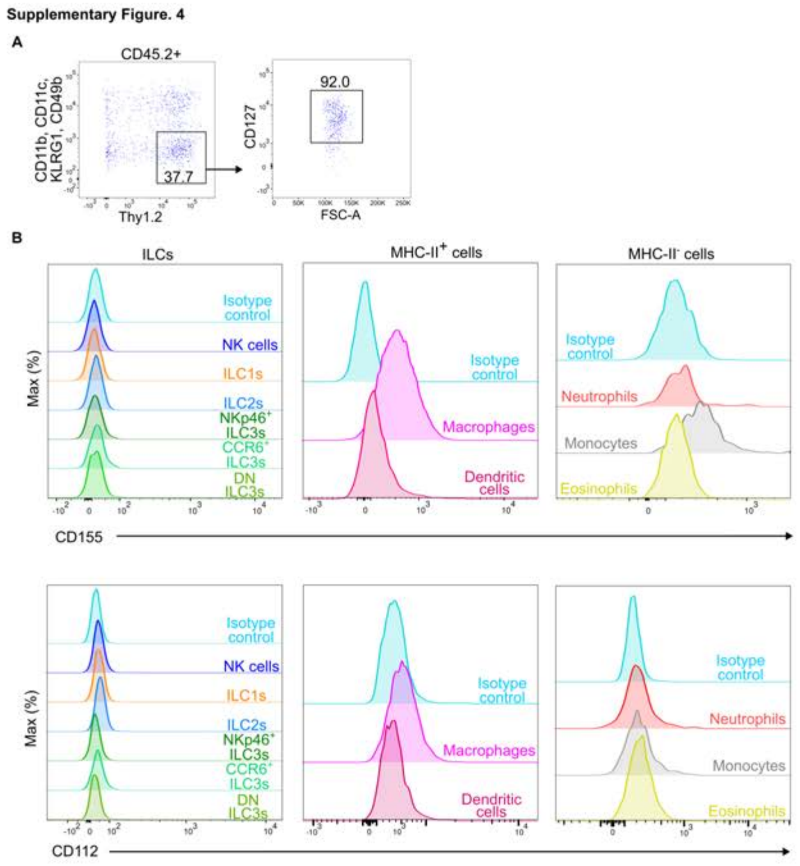


**Supplementary Figure 4.**

**(A)** Gating strategy for intestinal ILC3 purification used in Figures 4D–K. CD45.2$^{+}$ CD11b$^{-}$ CD11c$^{-}$ KLRG1$^{-}$ CD49b$^{-}$ Thy1.2$^{+}$ CD127$^{+}$ cells were sorted as ILC3s. **(B)** Representative histograms of CD155 (top) and CD112 (bottom) expression on intestinal immune cell populations from naive SCID mice. ILCs (NK cells, ILC1s, ILC2s, NKp46$^{+}$ ILC3s, CCR6$^{+}$ ILC3s, and DN ILC3s), MHC-II$^{+}$ cells (macrophages, dendritic cells), and MHC-II$^{-}$ cells (neutrophils, monocytes, eosinophils) were defined as described in Supplementary Figures 1A–C. Isotype control staining is shown for each group.

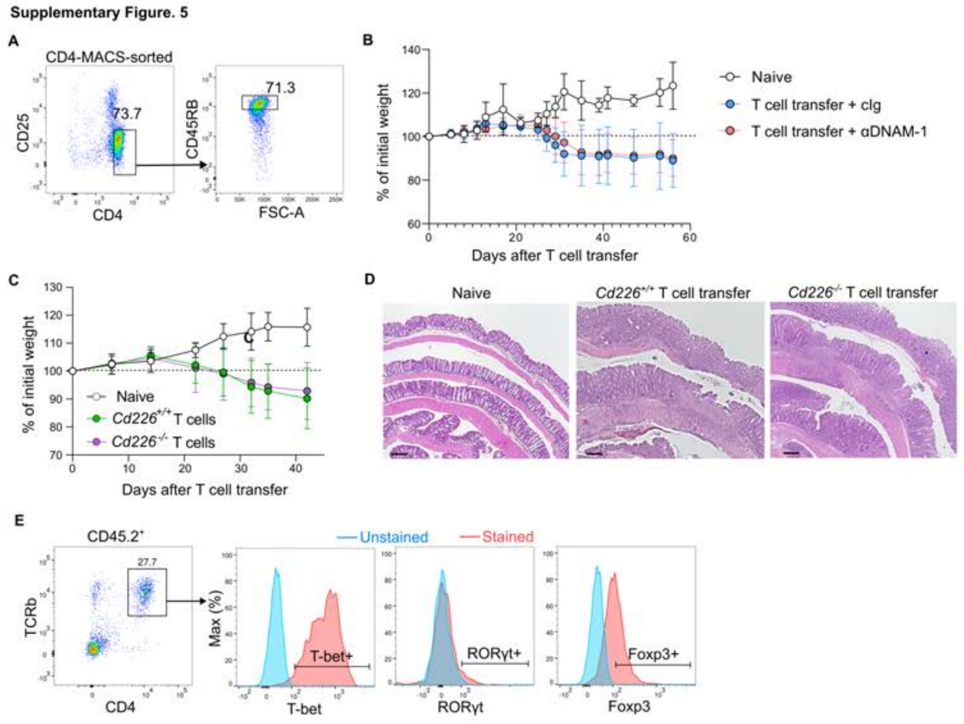


**Supplementary Figure 5.**

**(A)** Gating strategy for naive CD4⁺ T cells used in the T cell transfer colitis model. Splenic CD4⁺ cells were enriched by MACS sorting, and CD4⁺ CD25⁻ CD45RB$^{hi}$ cells were sorted by flow cytometry. **(B)** Body weight changes (% of initial weight) in naive mice (n = 3) and mice treated with cIg or anti–DNAM-1 mAb (n = 10) following T cell transfer (see Figures 5A–C). Data were pooled from two independent experiments. **(C)** Body weight changes (% of initial weight) in naive mice (n = 4) and mice transferred with WT or *Cd226*⁻/⁻ CD4⁺ T cells (n = 14–15) (see Figures 5D–F). Data were pooled from two independent experiments. **(D)** Representative H&E-stained colon sections from naive mice and mice transferred with WT or *Cd226*⁻/⁻ CD4⁺ T cells at day 49 after T cell transfer (see Figure 5F). Scale bars, 100 μm. **(E)** Gating strategy for CD4⁺ T cell subsets in mesenteric lymph nodes (see Figure 5J). CD45.2⁺ TCRβ⁺ CD4⁺ cells were gated, and T-bet⁺, RORγt⁺, and Foxp3⁺ populations were identified. Unstained controls are shown (blue) alongside stained samples (red). Data in (B, C) are shown as mean ± SD. Statistical significance was determined by two-way ANOVA followed by Šídák's multiple comparisons test.

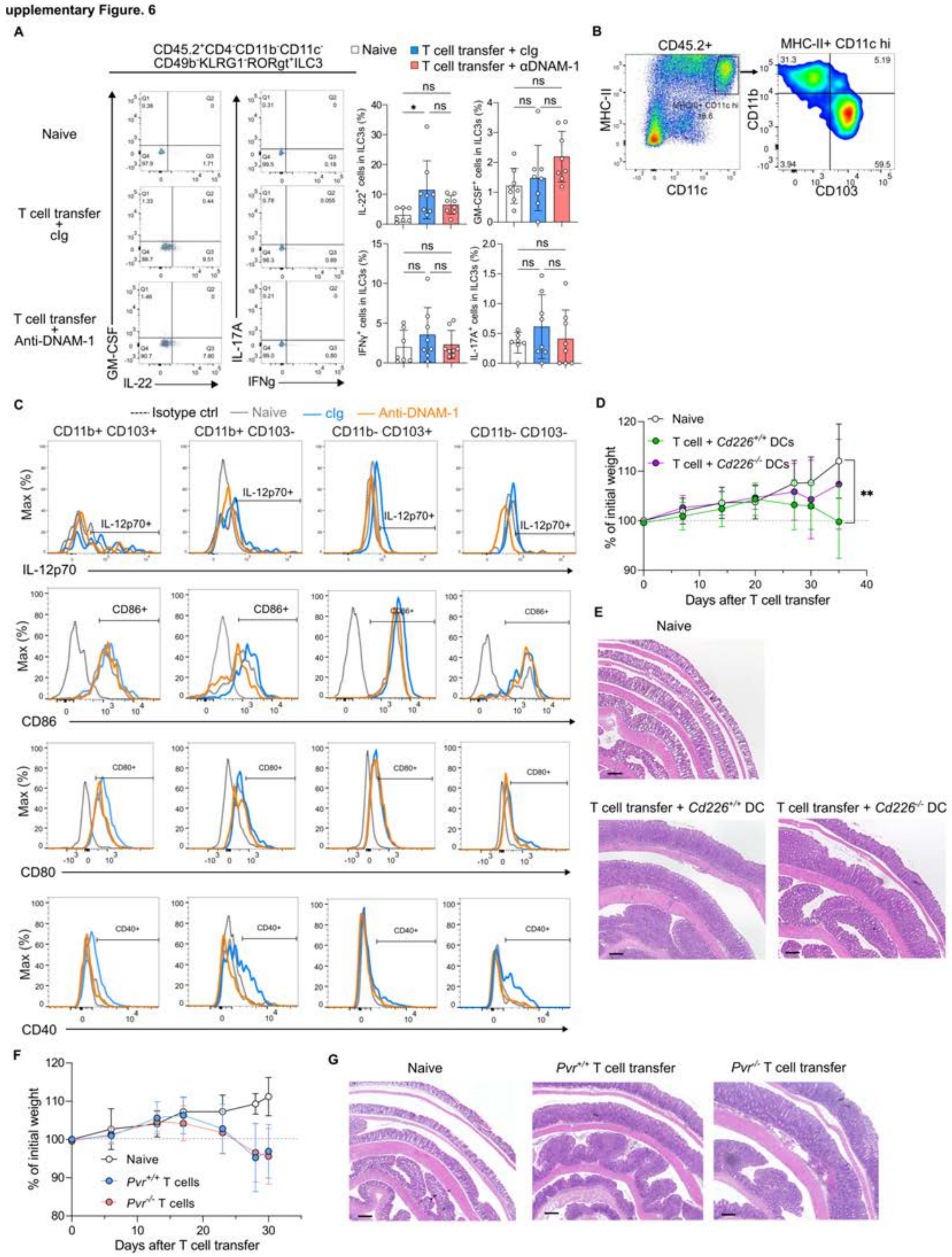


**Supplementary Figure 6.**

**(A)** Representative flow cytometry plots (left) and frequencies (right) of IL-22$^+$, GM-CSF$^+$, IFN-γ$^+$, and IL-17A$^+$ cells among intestinal ILC3s (CD45.2$^+$ CD4$^−$ CD11b$^−$ CD11c$^−$ CD49b$^−$ KLRG1$^−$ RORγt$^+$) from naive mice (n = 8) and mice treated with cIg or anti–

DNAM-1 mAb (n = 8) at day 10 after *Cd226*$^{-/-}$ CD4$^{+}$ T cell transfer. Data were pooled from two independent experiments. **(B)** Gating strategy for colonic DC subsets used in Figures 6A, B and Supplementary Figure 6C. CD45.2$^{+}$ MHC-II$^{+}$ CD11c$^{hi}$ cells were defined as DCs and further subdivided by CD11b and CD103 expression into four subsets (CD11b$^{+}$ CD103$^{+}$, CD11b$^{+}$ CD103$^{-}$, CD11b$^{-}$ CD103$^{+}$, and CD11b$^{-}$ CD103$^{-}$). **(C)** Representative histograms of IL-12p70, CD86, CD80, and CD40 expression on each colonic DC subset from naive mice (gray), colitic mice treated with cIg (blue), and colitic mice treated with anti–DNAM-1 mAb (orange) at day 7 after T cell transfer (see Figure 6B). Isotype control staining is shown as a dashed black line. **(D)** Body weight changes (% of initial weight) in naive mice (n = 10) and mice reconstituted with WT or Cd226$^{-}$/$^{-}$ DCs (n = 15–18) following *Cd226*$^{-/-}$ CD4$^{+}$ T cell transfer (see Figures 6C–E). Data were pooled from three independent experiments. **(E)** Representative H&E-stained colon sections from naive mice and mice reconstituted with WT or *Cd226*$^{-/-}$ DCs at day 35 after T cell transfer (see Figure 6E). Scale bars, 100 μm. **(F)** Body weight changes (% of initial weight) in naive mice (n = 9) and mice transferred with WT or *Pvr*$^{-/-}$ CD4$^{+}$ T cells (n = 8–10) (see Figures 6F, G). Data are representative of two independent experiments. **(G)** Representative H&E-stained colon sections from naive mice and mice transferred with WT or *Pvr*$^{-/-}$ CD4$^{+}$ T cells at day 30 after T cell transfer (see Figure 6G). Scale bars, 100 μm. Data in (A) are shown as mean ± SD. Statistical significance was determined by one-way ANOVA followed by Tukey's multiple comparisons test (A) or two-way ANOVA followed by Šídák's multiple comparisons test (D, F); **$p < 0.01$; ns, not significant.

Supplementary Figure. 7

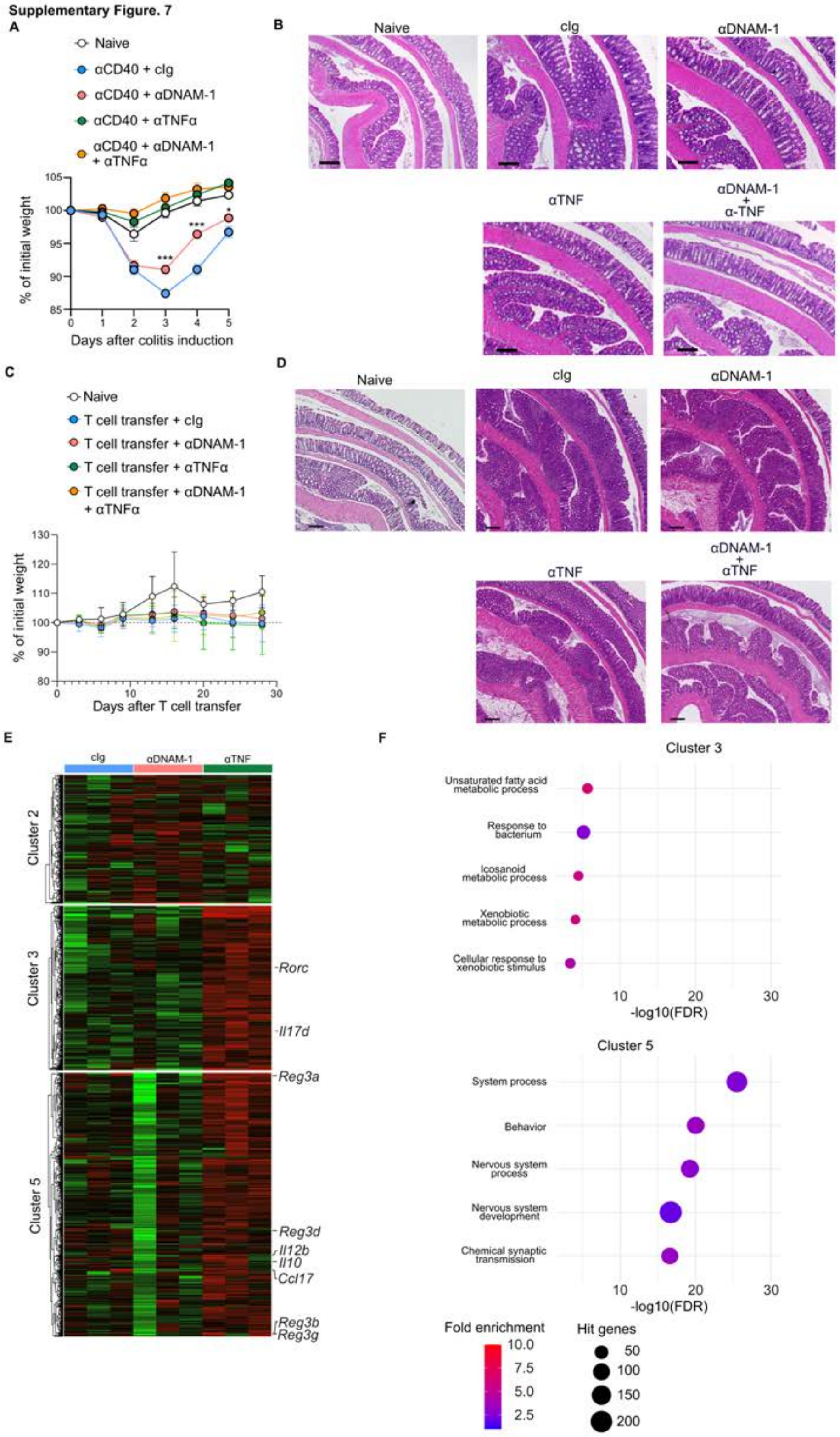

G

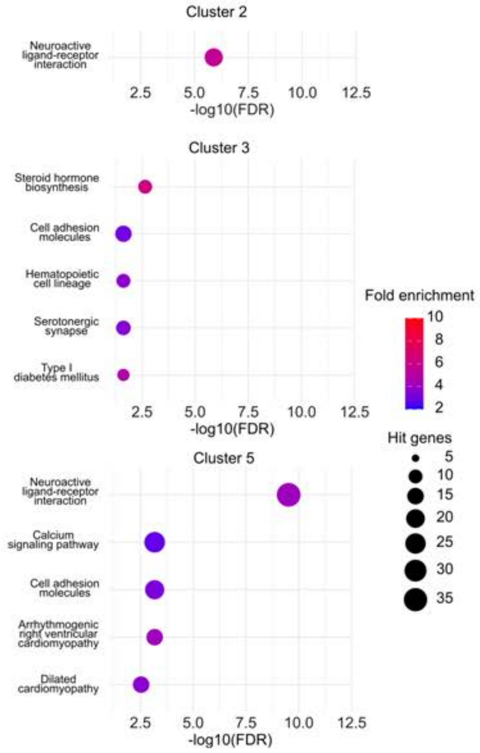


**Supplementary Figure 7.**

**(A)** Body weight changes (% of initial weight) in naive mice (n = 10) and mice treated with cIg, anti–DNAM-1 mAb, anti-TNFα mAb, or a combination of anti–DNAM-1 and anti-TNFα mAbs (n = 8–19) following anti-CD40 mAb injection (see Figure 7A). Data were pooled from three independent experiments. **(B)** Representative H&E-stained colon sections from naive mice and mice treated with cIg, anti–DNAM-1 mAb, anti-TNFα mAb, or a combination of anti–DNAM-1 and anti-TNFα mAbs at day 5 after anti-CD40 mAb injection (see Figure 7A). Scale bars, 100 μm. **(C)** Body weight changes (% of initial weight) in naive mice (n = 4) and mice treated with cIg, anti–DNAM-1 mAb, anti-TNFα mAb, or a combination of anti–DNAM-1 and anti-TNFα mAbs (twice weekly; n = 10–15) following T cell transfer (see Figures 7B). Data are representative of two independent experiments. **(D)** Representative H&E-stained colon sections from naive mice and mice treated with cIg, anti–DNAM-1 mAb, anti-TNFα mAb, or a combination of anti–DNAM-1 and anti-TNFα mAbs at day 28 after T cell transfer (see Figure 7B). Scale bars, 100 μm. **(E)** Heatmap of differentially expressed genes in Clusters 2, 3, and 5 identified by RNA sequencing of colonic tissues from mice treated with cIg, anti–DNAM-1 mAb, or anti-

TNFα mAb (n = 3 per group) at day 3 after anti-CD40 mAb injection (see Figure 7D). Representative genes are indicated. Color scale represents Z-score of normalized expression values. **(F)** Gene ontology (GO) biological process enrichment analysis of genes in Clusters 2, 3, and 5. Dot size represents the number of hit genes; color represents fold enrichment (see Figure 7E). **(G)** KEGG pathway enrichment analysis of genes in Clusters 2, 3, and 5. Dot size represents the number of hit genes; color represents fold enrichment (see Figure 7F). Data in (A, C) are shown as mean ± SD. Statistical significance was determined by two-way ANOVA followed by Šídák's multiple comparisons test; $^{*}p < 0.05$, $^{***}p < 0.001$.
